# Nonlinear transient amplification in recurrent neural networks with short-term plasticity

**DOI:** 10.1101/2021.06.09.447718

**Authors:** Yue Kris Wu, Friedemann Zenke

## Abstract

To rapidly process information, neural circuits have to amplify specific activity patterns transiently. How the brain performs this nonlinear operation remains elusive. Hebbian assemblies are one possibility whereby symmetric excitatory connections boost neuronal activity. However, such Hebbian amplification is often associated with dynamical slowing of network dynamics, non-transient attractor states, and pathological run-away activity. Feedback inhibition can alleviate these effects but typically linearizes responses and reduces amplification gain. At the same time, other alternative mechanisms rely on asymmetric connectivity, in conflict with the Hebbian doctrine. Here we propose nonlinear transient amplification (NTA), a plausible circuit mechanism that reconciles symmetric connectivity with rapid amplification while avoiding the above issues. NTA has two distinct temporal phases. Initially, positive feedback excitation selectively amplifies inputs that exceed a critical threshold. Subsequently, short-term plasticity quenches the run-away dynamics into an inhibition-stabilized network state. By characterizing NTA in supralinear network models, we establish that the resulting onset transients are stimulus selective and well-suited for speedy information processing. Further, we find that excitatory-inhibitory co-tuning widens the parameter regime in which NTA is possible. In summary, NTA provides a parsimonious explanation for how excitatory-inhibitory co-tuning and short-term plasticity collaborate in recurrent networks to achieve transient amplification.

## Introduction

Perception in the brain is reliable and strikingly fast. Recognizing a familiar face or locating an animal in a picture only takes a split second (Thorpe et al., 1996). This pace of processing is truly remarkable since it involves several recurrently connected brain areas each of which has to selectively amplify or suppress specific signals before propagating them further. This processing is mediated through circuits with several intriguing properties. First, excitatory-inhibitory (EI) currents into individual neurons are commonly correlated in time and co-tuned in stimulus space (Wehr and Zador, 2003; Froemke et al., 2007; Okun and Lampl, 2008; Hennequin et al., 2017; Rupprecht and Friedrich, 2018; Znamenskiy et al., 2018). Second, neural responses to stimulation are shaped through diverse forms of short-term plasticity (STP) (Tsodyks and Markram, 1997; Markram et al., 1998; Zucker and Regehr, 2002; Pala and Petersen, 2015). Finally, mounting evidence suggests that amplification rests on neuronal ensembles with strong recurrent excitation (Marshel et al., 2019; Peron et al., 2020), whereby excitatory neurons with similar tuning preferentially form reciprocal connections (Ko et al., 2011; Cossell et al., 2015). Such predominantly symmetric connectivity between excitatory cells is consistent with the notion of Hebbian cell assemblies (Hebb, 1949), which are considered an essential component of neural circuits and the putative basis of associative memory (Harris, 2005; Josselyn and Tonegawa, 2020). Computationally, Hebbian cell assemblies can amplify specific activity patterns through positive feedback, also referred to as Hebbian amplification. Based on these principles, several studies have shown that Hebbian amplification can drive persistent activity that outlasts a preceding stimulus (Hopfield, 1982; Amit and Brunel, 1997; Yakovlev et al., 1998; Wong and Wang, 2006; Zenke et al., 2015; Gillary et al., 2017), comparable to selective delay activity observed in the prefrontal cortex when animals are engaged in working memory tasks (Funahashi et al., 1989; Romo et al., 1999).

However, in most brain areas, evoked responses are transient and sensory neurons typically exhibit pronounced stimulus *onset* responses, after which the circuit dynamics settle into a low-activity steady-state even when the stimulus is still present (DeWeese et al., 2003; Mazor and Laurent, 2005; Bolding and Franks, 2018). Preventing run-away excitation and multi-stable attractor dynamics in recurrent networks requires powerful and often finely tuned feedback inhibition resulting in EI balance (Amit and Brunel, 1997; Compte et al., 2000; Litwin-Kumar and Doiron, 2012; Ponce-Alvarez et al., 2013; Mazzucato et al., 2019). However, feedback inhibition tends to linearize steady-state activity (Van Vreeswijk and Sompolinsky, 1996; Baker et al., 2020) and does not necessarily generate pronounced onset responses consistent with experiments. While feedforward inhibition provides one possible explanation for transient onset dynamics (Wehr and Zador, 2003; Vogels et al., 2011; Gjoni et al., 2018), it does not explain the recurrent excitation commonly seen in cortical circuits. As a possible remedy, balanced amplification constitutes a putative mechanism for transient amplification in recurrent neural networks (Murphy and Miller, 2009). However, to achieve strong amplification, several ensembles need to be chained together into a hidden feedforward structure which manifests in strongly non-normal recurrent connectivity (Goldman, 2009; Hennequin et al., 2012, 2014; Bondanelli and Ostojic, 2020; Gillett et al., 2020). Yet, such network structures are at odds with the often observed symmetric excitatory connectivity (Ko et al., 2011; Cossell et al., 2015).

We are thus faced with a conundrum. On the one hand, Hebbian assemblies, whose connectivity is consistent with neurobiology, can amplify specific stimuli. But, the resulting persistent attractor dynamics are inconsistent with the transient activity observed in experiments. On the other hand, non-normal connectivity offers an appealing explanation for transient amplification in recurrent neural network models, but it is at odds with the mainly observed symmetric connectivity. Importantly, however, previous studies largely ignored STP, which considerably modulates synaptic transmission and shapes neural responses on timescales ranging from milliseconds to minutes (Tsodyks and Markram, 1997; Markram et al., 1998; Zucker and Regehr, 2002; Pala and Petersen, 2015). This raises the question of whether and how STP or other neuronal adaptation mechanisms could resolve the puzzle by reconciling the seemingly disparate aspects.

Here we address this question by studying the emergence of transient dynamics in recurrent network models and examine how they are shaped through neuronal nonlinearities, STP, and EI cotuning. Specifically, we first characterize the conditions under which individual neuronal ensembles with symmetric excitatory connectivity succumb to explosive run-away activity in response to external stimulation. We then show how STP can effectively mitigate this instability by re-stabilizing ensemble dynamics in an inhibition-stabilized network (ISN) state, but only after generating a pronounced stimulus-triggered onset transient. We call this mechanism nonlinear transient amplification (NTA) and show that it yields selective onset responses that carry more relevant stimulus information than the subsequent steady-state. Finally, we characterize the functional benefits of global EI balance and co-tuning for NTA. We find that pattern classification in networks with NTA is enhanced by EI balance in individual neurons, a feature that is widely observed in the brain (Wehr and Zador, 2003; Froemke et al., 2007; Okun and Lampl, 2008; Rupprecht and Friedrich, 2018) and readily emerges in computational models endowed with activity-dependent plasticity of inhibitory synapses (Vogels et al., 2011). Importantly, NTA purports that, following transient amplification, neuronal ensembles settle into a stable ISN state, consistent with previous work on stabilized supralinear networks (SSNs) (Ahmadian et al., 2013; Rubin et al., 2015; Hennequin et al., 2018; Kraynyukova and Tchumatchenko, 2018). In summary, our work indicates that NTA is ideally suited to amplify stimuli rapidly through the interaction of symmetric recurrent excitation with STP.

## Results

To understand the emergence of transient responses in recurrent neural networks, we studied rate-based population models with a supralinear, power law input-output function (Fig. 1A, B; Ahmadian et al., 2013; Hennequin et al., 2018), which captures essential aspects of neuronal activation (Priebe et al., 2004), while also being analytically tractable. We first considered an isolated neuronal ensemble consisting of one excitatory (E) and one inhibitory (I) population (Fig. 1A).

**Fig. 1.**
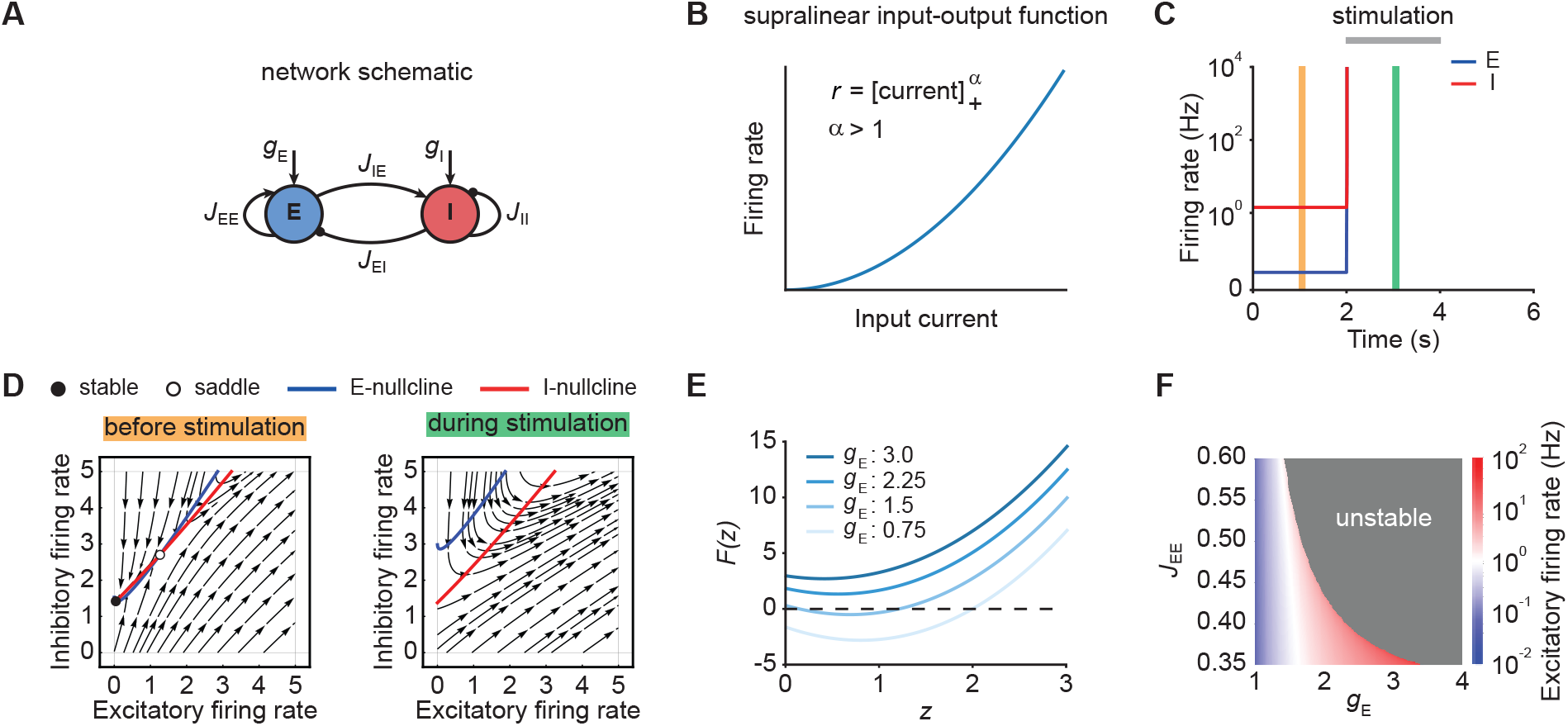
Neuronal ensembles nonlinearly amplify inputs above a critical threshold. **(A)** Schematic of the recurrent ensemble model consisting of an excitatory (blue) and an inhibitory population (red). **(B)** Supralinear input-output function given by a rectified power law with exponent *α* = 2. **(C)** Firing rates of the excitatory (blue) and inhibitory population (red) in response to external stimulation during the interval from 2–4s (gray bar). The stimulation was implemented by temporarily increasing the input *g_E_*. **(D)** Phase portrait of the system before stimulation (left; cf. C orange) and during stimulation (right; cf. C green). **(E)** Characteristic function *F*(*z*) for varying input strength *g_E_*. Note that the function loses its zero crossings, which correspond to fixed points of the system for increasing external input. **(F)** Heat map showing the evoked firing rate of the excitatory population for different parameter combinations *J_EE_* and *g_E_*. The gray region corresponds to the parameter regime with unstable dynamics.

The dynamics of this network are given by

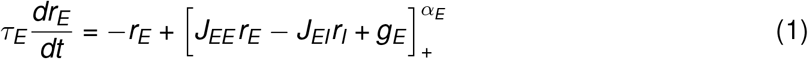

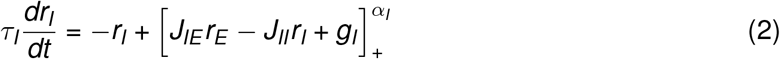

where *r_E_* and *r_I_* are the firing rates of the excitatory and inhibitory population, *τ_E_* and *τ_I_* represent the corresponding time constants, *J_XY_* denotes the synaptic strength from the population *Y* to the population *X*, where *X*, *Y* ∈ {*E*, *I*}, *g_E_* and *g_I_* are the external inputs to the respective populations. Finally, *α_E_* and *α_I_*, the exponents of the respective input-output functions, are fixed at two unless mentioned otherwise. For ease of notation, we further define the weight matrix **J** of the compound system as follows:

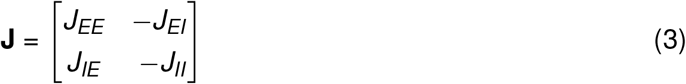

To account for the strong reciprocal E-to-E synaptic connections (Ko et al., 2011; Cossell et al., 2015), we studied networks in which the determinant

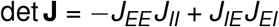

is negative. To mimic sensory stimulation, we investigated ensemble dynamics as a function of external input strength *g_E_*. Importantly, we assumed that most inhibition originates from recurrent connections and (Franks et al., 2011; Large et al., 2016), hence, we kept the input to the inhibitory population *g_I_* fixed.

### Nonlinear amplification of inputs above a critical threshold

We initialized the network in a stable low-activity state in the absence of external stimulation, consistent with spontaneous activity in cortical networks (Fig. 1C). However, an input *g_E_* of sufficient strength, destabilized the network (Fig. 1C). Importantly, this behavior is distinct from linear network models in which the network stability is independent of inputs (Methods). The transition from stable to unstable dynamics can be understood by examining the phase portrait of the system (Fig. 1D). Before stimulation, the system has a stable and an unstable fixed point (Fig. 1D, left). However, both fixed points disappear for an input *g_E_* above a critical stimulus strength (Fig. 1D, right).

To further understand the system’s bifurcation structure, we consider the characteristic function

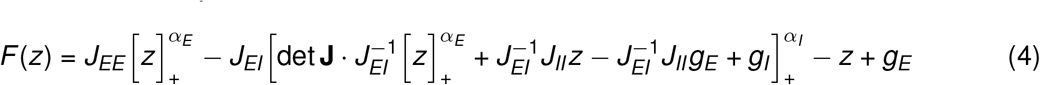

where *z* denotes the total current into the excitatory population and det **J** represents the determinant of the weight matrix (Kraynyukova and Tchumatchenko, 2018; Methods). The characteristic function reduces the original two-dimensional system to one dimension, whereby the zero crossings of the characteristic function correspond to the fixed points of the original system (cf. Eq. 1–2). We use this correspondence to visualize how the fixed points of the system change with the input *g_E_*. Increasing *g_E_* shifts *F*(*z*) upwards, which eventually leads to all zero crossings disappearing and the ensuing unstable dynamics (Fig. 1E; Methods). Importantly, for any weight matrix **J** with negative determinant, there exists a critical input *g_E_* at which all fixed points disappear (Methods). While for weak recurrent E-to-E connection strength *J_EE_*, the transition from stable dynamics to unstable is gradual, in that it happens at higher firing rates (Fig. 1F), it becomes more abrupt for stronger *J_EE_*. Thus, our analysis demonstrates that individual neuronal ensembles with negative determinant det **J** nonlinearly amplify inputs above a critical threshold by switching from initially stable to unstable dynamics.

### Short-term plasticity, but not spike-frequency adaptation, can re-stabilize ensemble dynamics

Since unstable dynamics are not observed in neurobiology, we wondered whether neuronal spike-frequency adaptation (SFA) or STP could re-stabilize the ensemble dynamics while keeping the nonlinear amplification character of the system. Specifically, we considered SFA of excitatory neurons, E-to-E short-term depression (STD), and E-to-I short-term facilitation (STF). We focused on these particular mechanisms because they are ubiquitously observed in the brain. Most pyramidal cells exhibit SFA (Barkai and Hasselmo, 1994) and most synapses show some form of STP (Markram et al., 1998; Zucker and Regehr, 2002; Pala and Petersen, 2015). Moreover, the time scales of these mechanisms are well-matched to typical timescales of perception, i.e., from milliseconds to seconds (Tsodyks and Markram, 1997; Fairhall et al., 2001; Pozzorini et al., 2013).

When we simulated our model with weak SFA, it did not stabilize run-away excitation (Fig. 2A; Methods). Increasing the strength of SFA eventually led to oscillatory ensemble activity (Fig. S1). To understand this behavior, we analyzed the corresponding characteristic function *F*(*z*) for an ensemble with SFA. We found that in the presence of SFA, *F*(*z*) can maximally have one stable low-activity fixed point and one unstable high-activity fixed point (Fig. S1). This property is closely related to SFA’s tendency to linearize a neuron’s input-output function (Ermentrout, 1998; Benda and Herz, 2003) but not to saturate it. Thus, when an input causes the system’s fixed points to disappear (cf. Fig. 1D, E), weak SFA either does not restore any fixed point, or the high-activity fixed point cannot “catch up” with the run-away dynamics. Therefore, the system’s dynamics remain unstable. Strong SFA, on the other hand, allows the high-activity fixed point to catch up with the increasing current into the excitatory population *z*, but since it is an unstable fixed point, this results in a reduction of the excitatory ensemble activity toward the low-activity fixed point. Unfortunately, this change is only short-lived and as the adaptation variable recovers (cf. Eq. (21)), the ensemble activity engages in another cycle of explosive run-away activity. Thus, while the input is present, strong SFA creates a stable limit cycle with associated oscillatory ensemble activity (Fig. S1, Methods), which was also shown in previous modeling studies (Van Vreeswijk and Hansel, 2001), but is not typically observed in neurobiology (DeWeese et al., 2003; Mazor and Laurent, 2005; Rupprecht and Friedrich, 2018).

**Fig. 2.**
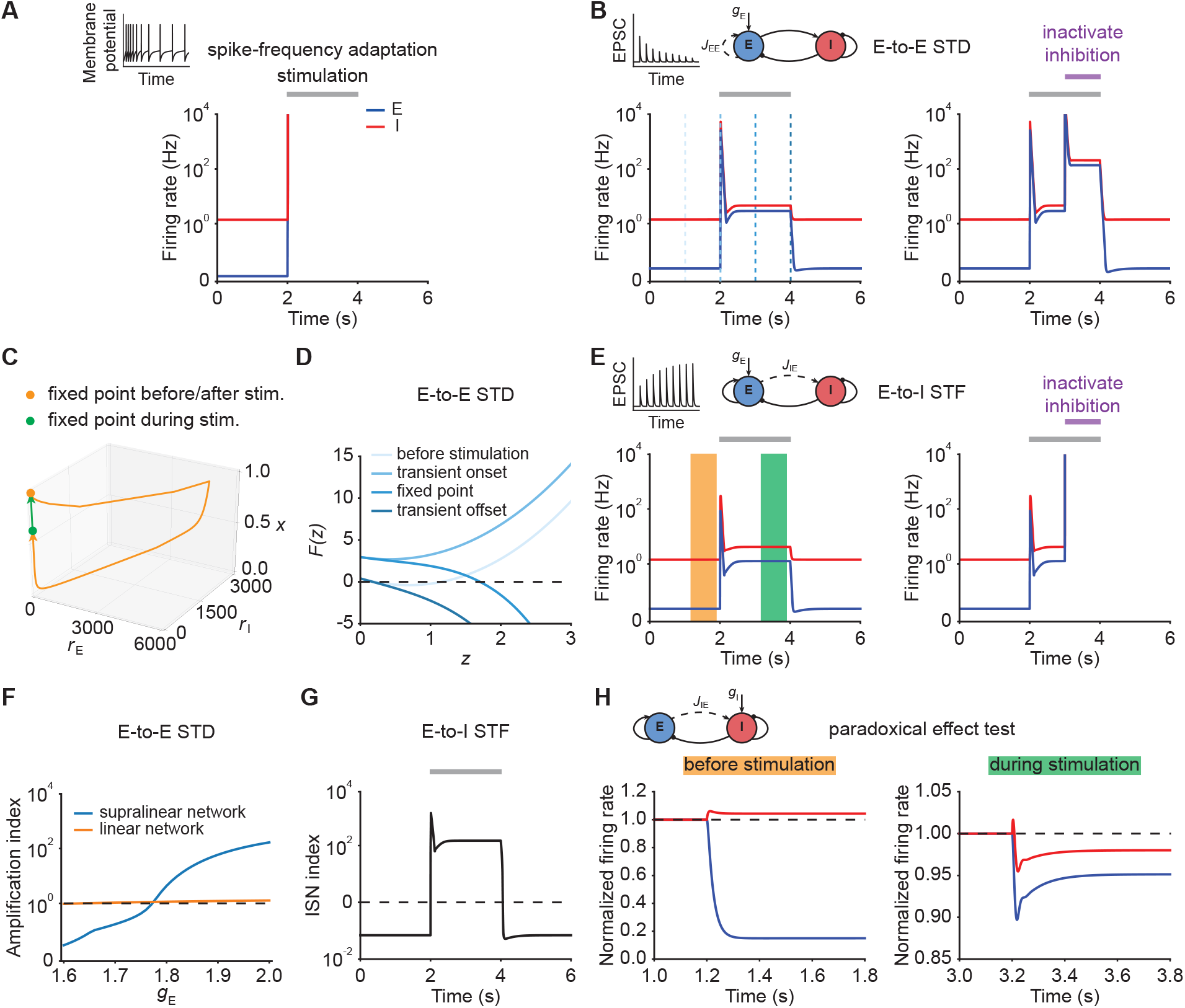
Short-term plasticity, but not spike-frequency adaptation, re-stabilizes ensemble dynamics. **(A)** Firing rates of the excitatory (blue) and inhibitory population (red) in the presence of spike-frequency adaptation (SFA). During stimulation (gray bar) additional input is injected into the excitatory population. The inset shows a cartoon of how SFA affects spiking neuronal dynamics in response to a step current input. **(B)** Left: Same as (A) but in the presence of E-to-E short-term depression (STD). Right: Same as left but inactivating inhibition in the period marked in purple. **(C)** 3D plot of the excitatory activity *r_E_*, inhibitory activity *r_I_*, and the STD variable *x* of the network in B left. The orange and green points mark the fixed points before/after and during stimulation. **(D)** Characteristic function *F*(*z*) in networks with E-to-E STD. Different brightness levels correspond to different time points in B left. **(E)** Same as (B) but in the presence of E-to-I short-term facilitation (STF). **(F)** Amplification index, which is defined as the ratio of the peak amplitude to input *g_E_*, as a function of input *g_E_* for supralinear networks (blue) and linear networks (orange) in the presence of E-to-E STD. **(G)** inhibition-stabilized network (ISN) index, which corresponds to the largest real part of the eigenvalues of the Jacobian matrix of the E-E subnetwork, as a function of time for the network with E-to-I STF in E left. For values above zero (dashed line), the ensemble is an ISN. **(H)** The normalized firing rates of the excitatory (blue) and inhibitory population (red) when injecting additional excitatory current into the inhibitory population before stimulation (left; cf. orange bar in E), and during stimulation (right; cf. green bar in E). Initially, the ensemble is in the non-ISN regime and injecting excitatory current into the inhibitory population increases its firing rate. During stimulation, however, the ensemble is an ISN. In this case, excitatory current injection into the inhibitory population results in a reduction of its firing rate, also known as the *paradoxical effect*.

Next, we considered STP, which is capable of saturating the effective neuronal input-output function (Mongillo et al., 2012; Zenke et al., 2015). We first analyzed the stimulus-evoked network dynamics when we added STD to the recurrent E-to-E connections. Strong depression of synaptic efficacy resulted in a brief onset transient after which the ensemble dynamics quickly settled into a stimulus-evoked steady-state with slightly higher activity than the baseline (Fig. 2B, left). After stimulus removal, the ensemble activity returned back to its baseline level (Fig. 2B, left; Fig. 2C). Notably, the ensemble dynamics remained stable, albeit at a much higher firing rate, when inhibition was inactivated during stimulus presentation (Fig. 2B, right). This shows that STP is capable of creating a stable high-activity fixed point, which is fundamentally different from the SFA dynamics discussed above. This difference in ensemble dynamics can be readily understood by analyzing the self-consistent solutions of *F*(*z*). Initially, the ensemble is at the stable low activity fixed point. But the stimulus causes this fixed point to disappear, thus giving way to positive feedback which creates the leading edge of the onset transient (cf. Fig. 2B). However, because E-to-E synaptic transmission is rapidly reduced by STD, the curvature of *F*(*z*) changes and a stable fixed point is created, thereby allowing excitatory run-away dynamics to terminate and the ensemble dynamics settle into a steady-state at low activity levels (Fig. 2D). We found that E-to-I STF leads to similar dynamics (Fig. 2E, left) with the only difference that this configuration requires inhibition for network stability (Fig. 2E, right), whereas E-to-E STD stabilizes activity even without inhibition, albeit at physiologically implausibly high activity levels. Importantly, the re-stabilization through either form of STP did not impair an ensemble’s ability to amplify stimuli during the initial onset phase.

To highlight the amplification power of supralinear networks over purely linear networks with equivalent weight strengths, we calculated the ratio of the evoked peak amplitudes of the firing rate to the input strength, henceforth called the “Amplification index”. Notably, amplification of stimuli above the critical threshold in supralinear networks is orders of magnitude larger than in a linear network with comparable weights (Fig. 2F). We stress that the resulting high firing rates are parameter-dependent (Fig. S2), but also due to the short duration of the onset peak. In experiments, such high rates are observed as precisely time-locked spikes (DeWeese et al., 2003; Wehr and Zador, 2003; Bolding and Franks, 2018; Gjoni et al., 2018).

Furthermore, we investigated how the network operating regime changes with the stimulation. Recent studies suggested that cortical networks operate as inhibition-stabilized networks (ISNs) (Sanzeni et al., 2020), in which the excitatory network is unstable in the absence of feedback inhibition (Tsodyks et al., 1997). Whether a network is an ISN or not is mathematically determined by the real part of the leading eigenvalue of the Jacobian of the excitatory-to-excitatory subnetwork (Tsodyks et al., 1997). We computed the leading eigenvalue in our model and referred to it as “ISN index” in the following (Methods). We found that the ISN index switches sign from negative to positive during external stimulation, indicating that the ensemble transitions from a non-ISN to an ISN (Fig. 2G). Notably, this behavior is distinct from linear network models in which the network operating regime is independent of the input (Methods). One defining characteristic of ISNs is that injecting excitatory (inhibitory) current into inhibitory neurons decreases (increases) inhibitory firing rates, which is also known as the paradoxical effect (Tsodyks et al., 1997; Miller and Palmigiano, 2020). To illustrate the difference in network operating regimes in terms of the paradoxical effect, we injected excitatory current into the inhibitory population before and during stimulus presentation. We found that before stimulation, the network did not exhibit the paradoxical effect (Fig. 2H, left; Fig. S3). In contrast, injecting excitatory inputs into the inhibitory population during stimulation reduced their activity (Fig. 2H, right; Fig. S3). Thus, during stimulation the neuronal ensemble switches to an ISN state.

Despite the fact that the supralinear input-output function of our framework captures some aspects of intracellular recordings (Priebe et al., 2004), it is unbounded and thus allows infinitely high firing rates. This is in contrast to neurobiology where firing rates are bounded due to neuronal refractory effects. While this assumption permitted us to analytically study the system and therefore to gain a deeper understanding of the underlying ensemble dynamics, we wondered whether our main conclusions were also valid when we limited the maximum firing rates. To that end, we carried out the same simulations while capping the firing rate at 300 Hz. In the absence of additional SFA or STP mechanisms, the firing rate saturation introduced a stable high-activity state in the ensemble dynamics which replaced the unstable dynamics in the uncapped model. As above, the ensemble entered this high-activity steady-state when stimulated with an external input above a critical threshold and exhibited persistent activity after stimulus removal (Fig. S4). While weak SFA did not change this behavior, strong SFA resulted in oscillatory behavior during stimulation consistent with previous analytical work (Fig. S4, Van Vreeswijk and Hansel, 2001), but did not in stable steady-states commonly observed in biological circuits. In the presence of E-to-E STD or E-to-I STF, however, the ensemble exhibited transient evoked activity at stimulation onset that was comparable to the uncapped case. Importantly, the ensemble did not show persistent activity after the stimulation (Fig. S4). Finally, we confirmed that all of these findings were qualitatively similar in a realistic spiking neural network model (Fig. S5; Methods).

In summary, we found that neuronal ensembles can rapidly, nonlinearly, and transiently amplify inputs by briefly switching from stable to unstable dynamics before being re-stabilized through STP mechanisms. We call this mechanism nonlinear transient amplification (NTA) which, in contrast to balanced amplification (Murphy and Miller, 2009; Hennequin et al., 2012), arises from nonlinear population dynamics interacting with STP. NTA is characterized by a large onset response, a subsequent ISN steady-state while the stimulus persists, and a return to a unique baseline activity state after the stimulus is removed. Thus, NTA is ideally suited to rapidly and nonlinearly amplify sensory inputs through symmetric recurrent excitatory connections, like reported experimentally (Ko et al., 2011; Cossell et al., 2015), while avoiding persistent activity.

### Co-tuned inhibition broadens the parameter regime of NTA

Up to now, we have focused on a single neuronal ensemble. However, to process information in the brain, several ensembles with different stimulus selectivity presumably coexist and interact in the same circuit. This coexistence creates potential problems. It can lead to multi-stable persistent attractor dynamics, which are not commonly observed and could have adverse effects on the processing of subsequent stimuli. One solution to this issue could be EI co-tuning, which arises in network models with plastic inhibitory synapses (Vogels et al., 2011) and has been observed experimentally in several sensory systems (Wehr and Zador, 2003; Froemke et al., 2007; Okun and Lampl, 2008; Rupprecht and Friedrich, 2018).

To characterize the conditions under which neuronal ensembles nonlinearly amplify stimuli without persistent activity, we analyzed the case of two interacting ensembles. More specifically, we considered networks with two excitatory ensembles and distinguished between global and co-tuned inhibition (Fig. 3A). In the case of global inhibition, one inhibitory population non-specifically inhibits both excitatory populations (Fig. 3A, left). In contrast, in networks with co-tuned inhibition, each ensemble is formed by a dedicated pair of an excitatory and an inhibitory population which can have cross-over connections, for instance, due to overlapping ensembles (Fig. 3A, right).

**Fig. 3.**
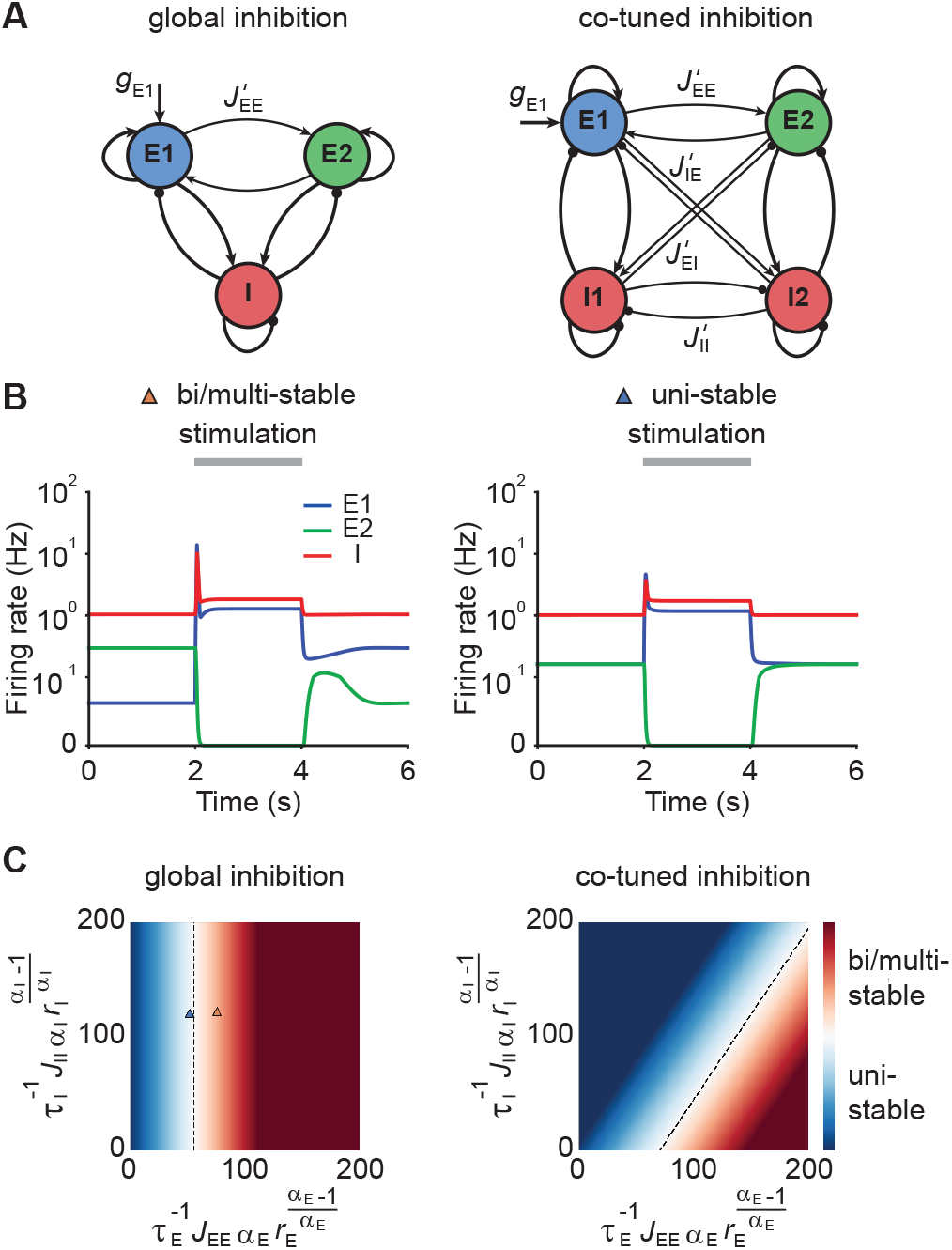
Co-tuned inhibition broadens the parameter regime of NTA. **(A)** Schematic of two neuronal ensembles with global inhibition (left) and with co-tuned inhibition (right). **(B)** Firing rate dynamics of bi/multi-stable ensemble dynamics (left) and uni-stable (right). In both cases, additional excitatory inputs are injected into excitatory ensemble *E*1 during the period marked in gray. **(C)** Analytical solution of uni- and bi/multi-stability regions for global inhibition (left) and co-tuned inhibition (right). Co-tuning results in a larger parameter regime of uni-stability. The triangles correspond to the two examples in B.

Global inhibition supports winner-take-all competition and is therefore often associated with multi-stable attractor dynamics (Wong and Wang, 2006; Mongillo et al., 2008). We first illustrated this effect in a network model with global inhibition. When the recurrent excitatory connections within each ensemble were sufficiently strong, small amounts of noise in the initial condition led to one of the ensembles spontaneously activating at elevated firing rates, while the other ensemble’s activity remained low (Fig. 3B, left). A specific external stimulation could trigger a switch from one state to the other in which the other ensemble was active at a high firing rate. Importantly, this change persisted even after the stimulus had been removed, a hallmark of multi-stable dynamics. In contrast, uni-stable systems have a global symmetric state in which both ensembles have the same activity in the absence of stimulation. While the stimulated ensemble showed elevated firing rates in response to the stimulus, its activity returned to the baseline level after the stimulus is removed (Fig. 3B, right), consistent with experimental observations (DeWeese et al., 2003; Rupprecht and Friedrich, 2018; Bolding and Franks, 2018). Note that the only difference between these two models is that *J_EE_* is larger in the multi-stable example than in the uni-stable one.

Symmetric baseline activity is most consistent with activity observed in sensory areas. Hence, we sought to understand which inhibitory connectivity would be most conducive to maintain it. To that end, we analytically identified the uni-stability conditions, which are determined by the leading eigenvalue of the Jacobian matrix of the system, for networks with varying degrees of EI co-tuning (Methods). We found that a broader parameter regime underlies uni-stability in networks with cotuned inhibition than global inhibition (Fig. 3C). Notably, this conclusion is general and extends to networks with an arbitrary number of ensembles (Methods). However, co-tuning does not impair an ensemble’s ability to exhibit NTA as shown above for an isolated ensemble with *perfect* cotuning. Thus, co-tuned inhibition helps to avoid persistent attractor dynamics and broadens the parameter regime of uni-stability without adversely affecting NTA.

### NTA provides better pattern completion and pattern separation than fixed points

Neural circuits are capable of generating stereotypical activity patterns in response to partial cues and forming distinct representations in response to different stimuli. To test whether NTA achieves pattern completion and supports pattern separation, we analyzed the transient onset activity in our models and compared it to the fixed point activity.

To investigate pattern completion and pattern separation in our model, we considered a co-tuned network with E-to-E STD and two distinct excitatory ensembles *E*1 and *E*2. We gave additional input *g*_*E*1_ to a Subset 1, consisting of 75% of the neurons in ensemble *E*1 (Fig. 4A). We then measured the evoked activity in the remaining 25% of the excitatory neurons in *E*1 to quantify pattern completion. To assess pattern separation, we injected additional input *g*_*E*1_ into the *E*1 neurons during the second stimulation phase (Fig. 4A) while measuring the activity of *E*2. Interestingly, we found that neurons in Subset 2, which did not receive additional input, showed large onset responses, their steady-state activity was largely suppressed (Fig. 4B). Despite the fact that inputs to *E*1 caused increased transient onset responses in *E*2, the amount of increase was orders of magnitude smaller than in *E*1 (Fig. 4B). To quantify pattern completion, we defined the

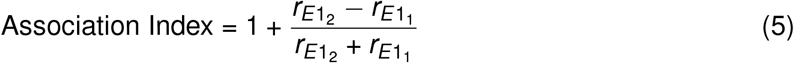

**Fig. 4.**
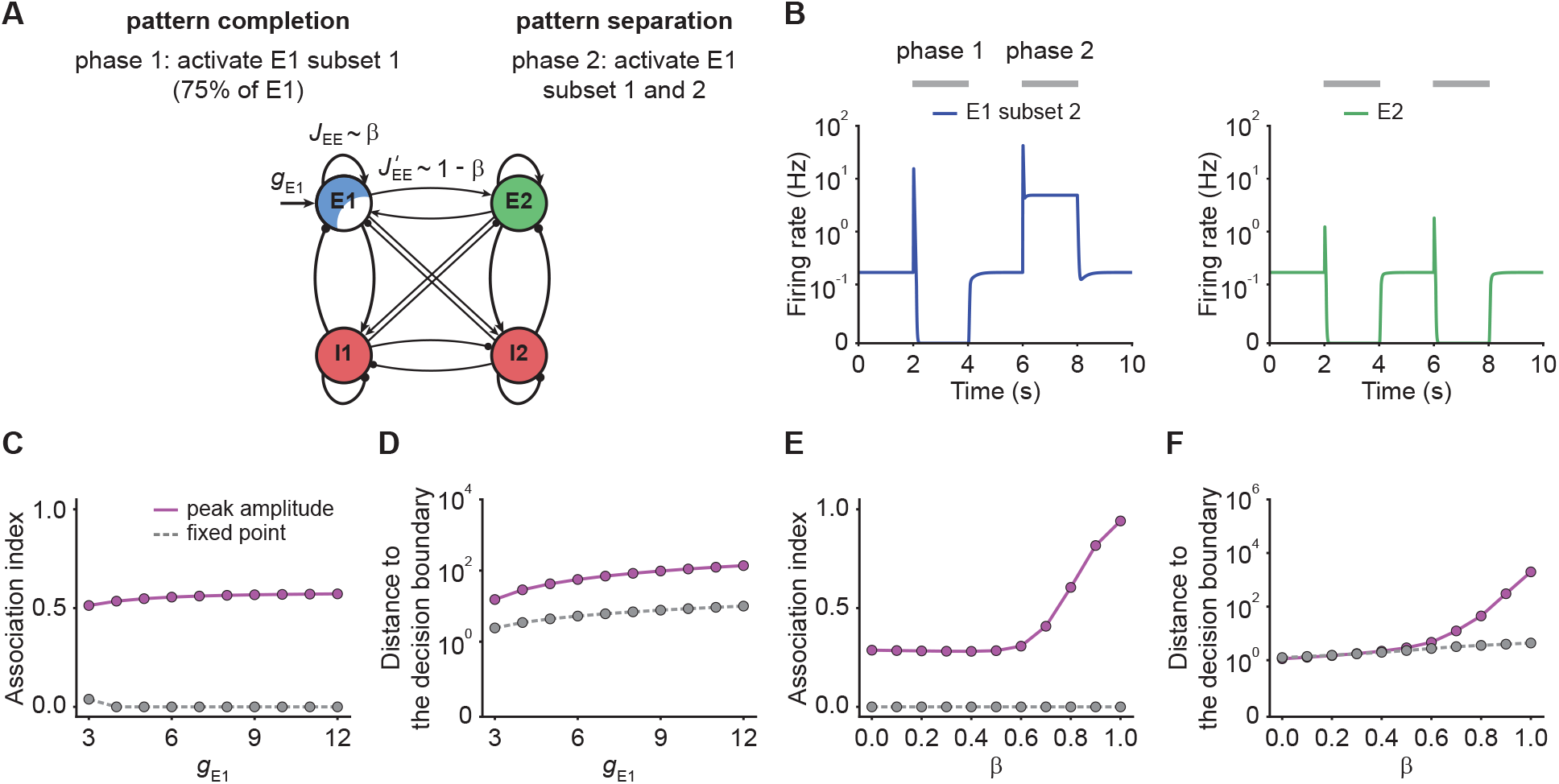
NTA yields stronger pattern completion and pattern separation than fixed points. **(A)** Schematic of the network setup used to probe pattern completion and pattern separation. To assess the effect on pattern completion, 75% of the neurons (Subset 1) in ensemble *E*1 received additional input *g*_*E*1_ during Phase 1 (2-4 s), while we recorded the firing rate of the remaining 25% (Subset 2) in the excitatory ensemble *E*1. To evaluate the impact on pattern separation, all neurons in *E*1 received additional inputs *g*_*E*1_ in Phase 2 (6-8 s) while the firing rate of *E*2 was measured. **(B)** Examples of firing rates of Subset 2 of *E*1 (left, blue) and *E*2 (right, green) with E-to-E STD. **(C)** Association index as a function of input *g*_*E*1_ for the onset peak amplitude (magenta solid line) and fixed-point activity (gray dashed line) for E-to-E STD. **(D)** Distance to the decision boundary as a function of input *g*_*E*1_ for the onset peak amplitude (magenta solid line) and fixed-point activity (gray dashed line) for E-to-E STD. **(E and F)** Same as C and D but as a function of *β*, which controls the inner- and inter-ensemble connection strength.

Here, *r*_*E*1_1__ and *r*_*E*1_2__ correspond to the subpopulation activities of *E*1, respectively. As per our definition, the Association Index ranges from zero to one, with larger values indicating stronger associativity. In addition, to quantify the separation between *E*1 and *E*2, we considered a binary classifier tasked to distinguish the two input stimuli and measured the distance to the classifier’s decision boundary, whereby larger values indicate a larger classification margin and thus better separability (Methods). Note that the Association Index and the distance to the decision boundary were computed from different input configurations corresponding to different phases in our simulation paradigm (Fig. 4B).

With these definitions, we ran simulations with different input strengths *g*_*E*1_. We found that the onset peaks showed stronger association than the fixed-point activity (Fig. 4C). Note that the association index at the fixed point remained zero, a direct consequence of *r*_*E*1_2__ being suppressed to zero (Fig. 4C). Furthermore, we found that the separation between the transient onset response and the decision boundary was always greater than for the fixed-point activity (Fig. 4D) showing that onset responses provide better pattern separation than fixed points.

To investigate how the recurrent excitatory connectivity affects both pattern completion and pattern separation, we introduced the parameter *β* which controls the within-ensemble E-to-E strength *J_EE_* relative to the inter-ensemble strength 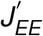 (Fig. 4A) such that *J_EE_* = *βJ*_tot_ and 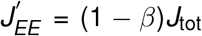. These definitions ensure that the total weight 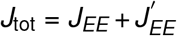 remains constant for any choice of *β*. Notably, the overall recurrent excitation strength within an ensemble *J_EE_* increases with increasing *β*. When *β* is larger than 0.5, the excitatory connection strength within the ensemble *J_EE_* exceeds the one between ensembles 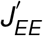.

We found that NTA’s pattern completion and separation capabilities monotonically increase with *β* (Fig. 4E, F), confirming that recurrent excitatory strength is a key determinant of network dynamics. Finally, we confirmed that our findings were also valid in networks with E-to-I STF (Fig. S6), which is commonly observed in the brain (Markram et al., 1998; Zucker and Regehr, 2002; Pala and Petersen, 2015). In summary, NTA’s transient onset responses result in overall better pattern completion and pattern separation than fixed point activity.

### NTA provides higher amplification and pattern separation in morphing experiments

So far, we only considered input to one ensemble. To examine how representations in our model are affected by ambiguous inputs to several ensembles, we performed additional morphing experiments (Freedman et al., 2001; Niessing and Friedrich, 2010). To that end, we introduced the parameter *p* which interpolates between two input stimuli which target *E*1 and *E*2 respectively. When *p* is zero, all additional input is injected into *E*1. For *p* equal to one, all additional input is injected into *E*2. Finally, *p* equal to 0.5 corresponds to the symmetric case in which *E*1 and *E*2 receive the same amount of additional input (Fig. 5A).

**Fig. 5.**
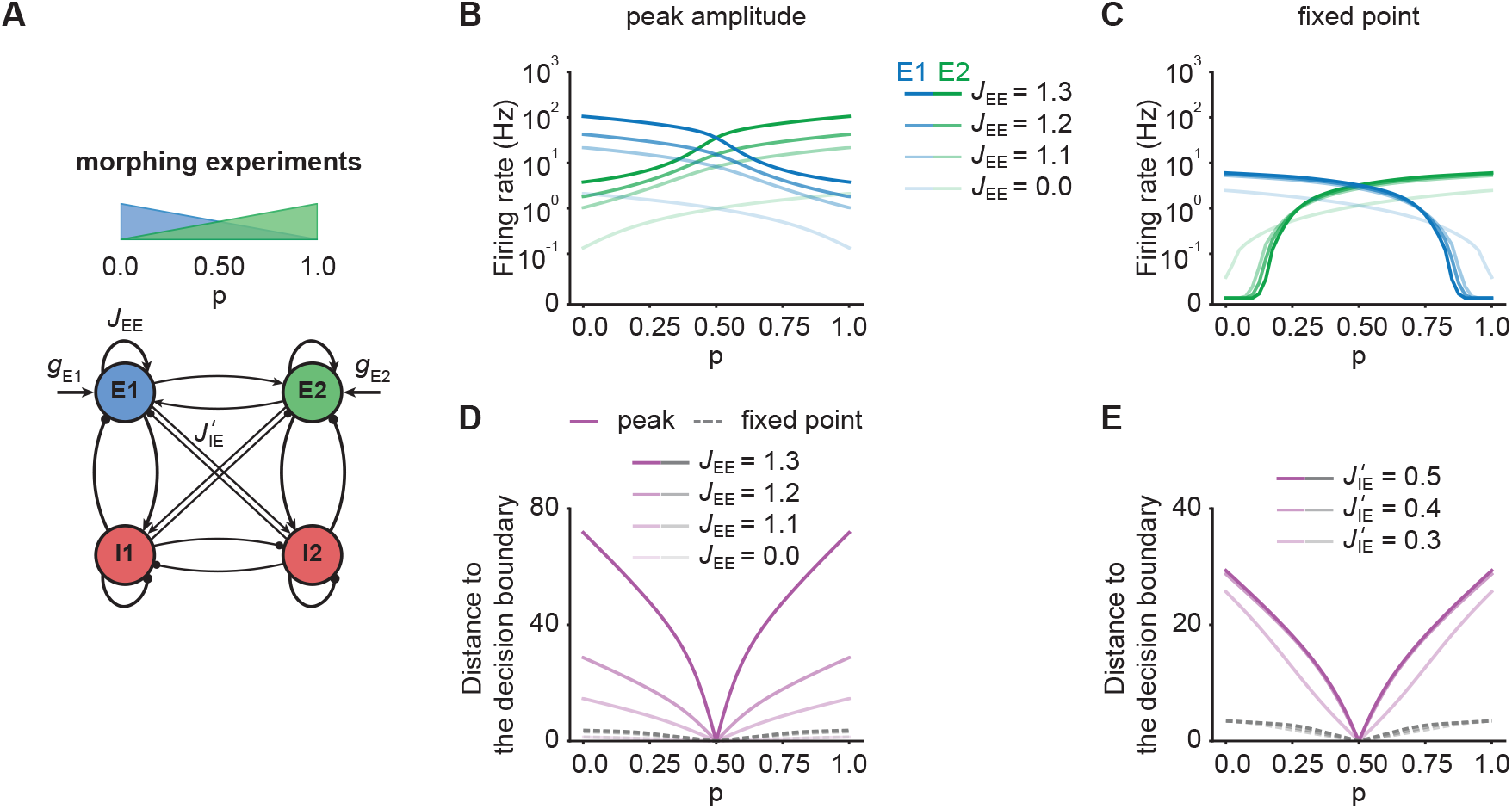
NTA provides stronger amplification and pattern separation in morphing experiments than fixed point activity. **(A)** Schematic of the morphing stimulation paradigm. The fraction of the additional inputs into the two excitatory ensembles is controlled by the parameter *p*. **(B)** Peak amplitude of *E*1 (blue) and *E*2 (green) as a function of *p*. Brightness levels represent different recurrent E-to-E connection strengths *J_EE_*. **(C)** Same as in B but for fixed-point activity. **(D)** Distance to the decision boundary as a function of *p* for the peak onset response (magenta solid line) and fixed-point activity (gray dashed line). **(E)** Same as D but with different E-to-I connection strengths 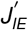 across ensembles.

First, we investigated how the recurrent excitatory connection strength within each ensemble *J_EE_* affects the onset peak amplitude and fixed-point activity. We found that the peak amplitudes depend strongly on *J_EE_*, whereas the fixed-point activity was only weakly dependent on *J_EE_* (Fig. 5B, C). When we disconnected the ensembles by completely eliminating all recurrent excitatory connections, activity was noticeably decreased (Fig. 5B, C). This illustrates, that recurrent excitation does play an important role in selectively amplifying specific stimuli similar to experimental observations (Marshel et al., 2019; Peron et al., 2020), but that amplification is highest at the onset.

Further, we examined the impact of competition through lateral inhibition as a function of the E-to-I inter-ensemble strength 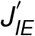 (Methods). As above, we quantified its impact by measuring the representational distance to the decision boundary for the transient onset responses and fixed-point activity. We found that regardless of the specific STP mechanism, the distance was larger for the onset responses than for the fixed-point activity, consistent with the notion that the onset can encode the stimulus identity more reliably than the fixed-point (Fig. 5D-E, Fig. S7). Thus, NTA provides stronger amplification and pattern separation than fixed points in response to ambiguous stimuli.

### Nonlinear transient amplification in spiking neural networks

Thus far, our analysis relied on power law neuronal input-output functions in the interest of analytical tractability. To test whether our findings also qualitatively apply to more realistic network models, we built a spiking neural network consisting of randomly connected 800 excitatory and 200 inhibitory neurons, in which the E-to-E synaptic connections were subject to STD (Methods). Here, we defined five overlapping ensembles, each corresponding to 200 randomly selected excitatory neurons. During an initial simulation phase (0–22s), we consecutively stimulated each pattern by giving additional input to their excitatory neurons, whereas the input to other neurons remained unchanged (Fig. 6A). In addition, we also tested pattern completion by stimulating only 75% (Subset 1) of the neurons belonging to Pattern 5 (22–24 s; Fig. 6A). We quantified each pattern’s activity by calculating the population firing rate of the stored patterns (Methods). As in the case of the rate-based model, the neuronal ensembles in the spiking model generated pronounced transient onset responses. We then measured the difference of peak pattern activity and steadystate activity between the stimulated pattern and the remaining unstimulated patterns (Methods). As for the rate-based networks, this difference was consistently larger for the onset peak than for the fixed-point (Fig. 6B, C). Thus, transient onset responses allow better stimulus separation than fixed points also in spiking neural network models.

**Fig. 6.**
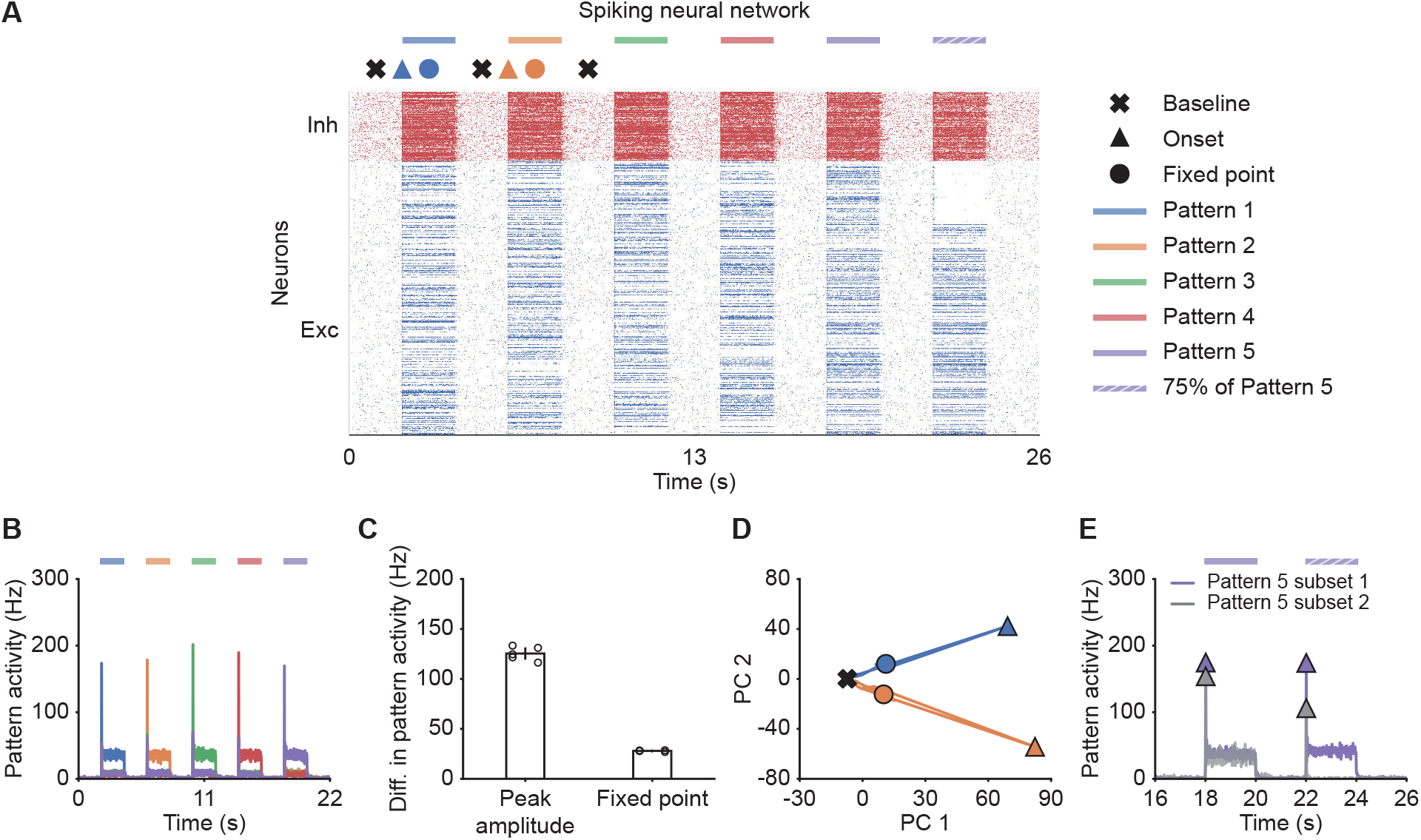
Spiking neural network simulations qualitatively reproduce NTA dynamics of rate models. **(A)** Spiking activity of excitatory (blue) and inhibitory (red) neurons in a spiking neural network. From 2–20s, Patterns 1–5 individually received additional input for 2 s each (colored bars). From 22–24 s, 75% of Pattern 5 neurons (Subset 1) received additional input, whereas the rest 25% of Pattern 5 neurons (Subset 2) did not receive additional input. The symbols at the top designate the different simulation phases of baseline activity, the onset transients, and the fixed point activity. Different colors correspond to the distinct stimulation periods. **(B)** Pattern activity of each stored pattern (colors). **(C)** Difference in pattern activity between the stimulated pattern with the remaining patterns for the transient onset peak and the fixed point. Points correspond to the different stimulation periods. **(D)** Spiking activity during the interval 0–10s represented in the PCA basis spanned by the first two principal components which captured approximately 40% of the total variance. The colored lines represent the PC trajectories of the first two stimuli shown in A and B. Triangles, points and crosses correspond to the onset peak, fixed point, and baseline activity, respectively. **(E)** Pattern activity of Subset 1 (purple) and Subset 2 (gray) of Pattern five from 16–26s. Onset peaks are marked by triangles.

Finally, to visualize the neural activity, we projected the binned spiking activity during the first 10 s of our simulation onto its first two principal components. Notably, the PC trajectory does not exhibit a pronounced rotational component (Fig. 6D) as activity is confined to one specific ensemble, consistent with experiments (Marshel et al., 2019). Furthermore, we computed the fifth pattern’s activity for Subset 1 and 2 during the time interval 16–26s. In agreement with our rate models, neurons in Subset 2 which did not receive additional inputs showed a strong response at the onset (Fig. 6E), but not at the fixed point, suggesting that the strongest pattern completion occurs during the initial amplification phase. Thus, the key characteristics of NTA are preserved across rate-based and more realistic spiking neural network models.

## Discussion

In this study, we demonstrated that neuronal ensemble models with recurrent excitation and suitable forms of STP exhibit nonlinear transient amplification (NTA), a putative mechanism underlying selective amplification in recurrent circuits. NTA combines a supralinear neuronal transfer function, recurrent excitation between neurons with similar tuning, and pronounced STP. Using analytical and numerical methods, we showed that NTA generates rapid transient onset responses during which optimal stimulus separation occurs rather than at steady-states. Additionally, we showed that co-tuned inhibition is conducive to prevent the emergence of persistent activity, which could otherwise interfere with processing subsequent stimuli. In contrast to balanced amplification (Murphy and Miller, 2009), NTA is an intrinsically nonlinear mechanism for which only stimuli above a critical threshold are amplified effectively. While the precise threshold value is parameter-dependent, it can be arbitrarily low provided the excitatory recurrent connections are sufficiently strong (cf. Fig. 1F). Importantly, such a critical activation threshold offers a possible explanation for sensory perception experiments which show similar threshold behavior (Marshel et al., 2019; Peron et al., 2020). Following transient amplification, ensemble dynamics are inhibition-stabilized, which renders our model compatible with existing work on SSNs (Ahmadian et al., 2013; Rubin et al., 2015; Hennequin et al., 2018; Kraynyukova and Tchumatchenko, 2018). Thus, NTA provides a parsimonious explanation for why sensory systems may rely upon neuronal ensembles with recurrent excitation in combination with EI co-tuning, and pronounced STP dynamics.

Several theoretical studies approached the problem of transient amplification in recurrent neural network models. One particularly well-studied mechanism relies on non-normal connectivity matrices whereby stimuli are selectively amplified through asymmetric synaptic connections with an implicit feedforward structure (Murphy and Miller, 2009; Goldman, 2009; Hennequin et al., 2014; Bondanelli and Ostojic, 2020; Gillett et al., 2020; Christodoulou et al., 2021). Importantly, nonnormal amplification can generate rich transient activity in linear network models but lacks a critical threshold above which amplification occurs. These properties contrast with NTA, which relies on a nonlinear transfer function and symmetric excitatory connections within neuronal ensembles with similar tuning. The symmetric connectivity generates potent run-away activity above a critical stimulus strength. Nevertheless, the overall network dynamics are stable because run-away dynamics are eventually quenched through STP. Crucially, after the transient amplification phase, ensemble dynamics settle in an inhibitory-stabilized state, which renders NTA compatible with previous work on SSNs only that in our case stabilization is accomplished dynamically through STP. Due to the switch of the network’s dynamical state, NTA’s amplification is orders of magnitudes larger than balanced amplification (Murphy and Miller 2009; cf. Fig. 2D, Fig. S8).

NTA requires STP and recurrent inhibition, whose role on network dynamics have been studied in the past. Yet, STP was mainly examined in the context of stable working memory in recurrent networks (Mongillo et al., 2008; Zenke et al., 2015; Seeholzer et al., 2019) and transient delay activity following stimulus offset (Hempel et al., 2000; Gillary et al., 2017). However, its role in generating strongly amplified onset transients as a possible coding paradigm was largely ignored. Recurrent inhibition is essential for NTA to ensure uni-stability and selectivity through the suppression of ensembles with different tuning. This requirement is similar in flavor to semi-balanced networks characterized by excess inhibition to some excitatory ensembles while others are balanced (Baker et al., 2020). However, the theory of semi-balanced networks has, so far, only been applied to steady-state dynamics while ignoring transients and STP. Previous work showed that STP can tune networks to a critical state (Levina et al., 2007), but focused primarily on E-to-E STD. Our work extends this notion by combining it with a distinct computational mechanism and shows that both E-to-E STD and E-to-I STF can re-stabilize ensemble dynamics. EI co-tuning prominently features in several models and was shown to support network stability (Vogels et al., 2011; Hennequin et al., 2017; Znamenskiy et al., 2018), efficient coding (Denève and Machens, 2016), novelty detection (Schulz et al., 2020), changes in neuronal variability (Hennequin et al., 2018; Rost et al., 2018), and correlation structure (Wu et al., 2020). Moreover, some studies have argued that EI balance and co-tuning could increase robustness to noise in the brain (Rubin et al., 2017). The present work mainly highlights its importance for preventing multi-stability and delay activity in circuits not requiring such long-timescale dynamics.

NTA is consistent with several experimental findings. First, our model recapitulates the key findings of Shew et al. (2015) who showed *ex vivo* that strong sensory inputs cause a transient shift to a supercritical state, after which adaptive changes rapidly tune the network to criticality. Second, NTA requires symmetric excitatory connectivity between neurons with similar tuning, which has been reported in experiments (Ko et al., 2011; Cossell et al., 2015; Peron et al., 2020). Third, ensemble activation in our model depends on a critical stimulus strength in line with recent all-optical experiments in the visual cortex, which further link ensemble activation with a perceptual threshold (Marshel et al., 2019). Fourth, sensory networks are uni-stable in that they return to a non-selective activity state after the removal of the stimulus and usually do not show persistent activity (DeWeese et al., 2003; Mazor and Laurent, 2005; Rupprecht and Friedrich, 2018). Fifth, our work shows that NTA’s onset responses encode stimulus identity better than the fixed-point activity, consistent with experiments in the locust antennal lobe (Mazor and Laurent, 2005) and research supporting that the brain relies on coactivity on short timescales to represent information (Stopfer et al., 1997; Engel et al., 2001; Harris et al., 2003; El-Gaby et al., 2021). Finally, EI co-tuning, which is conducive for NTA, has been found ubiquitously in different sensory circuits (Wehr and Zador, 2003; Froemke et al., 2007; Okun and Lampl, 2008; Rupprecht and Friedrich, 2018; Znamenskiy et al., 2018).

In our model, we made several simplifying assumptions. For instance, we kept the input to inhibitory neurons fixed and only varied the input to the excitatory population. This step was motivated by experiments in the piriform cortex where the total inhibition is dominated by feedback inhibition (Franks et al., 2011). Nevertheless, significant feedforward inhibition was observed in other areas (Bissieére et al., 2003; Cruikshank et al., 2007; Ji et al., 2016; Miska et al., 2018). While an in-depth comparison for different origins of inhibition was beyond the scope of the present study, we found that increasing the inputs to the excitatory population and inhibitory population by the same amount can still lead to NTA (Fig. S9; Methods). Therefore, we are confident that our main findings remain unaffected in the presence of substantial feedforward inhibition. Similarly, we limited our analysis to only a few overlapping ensembles. It will be interesting future work to study NTA in the case of many interacting and potentially overlapping ensembles and to determine the maximum storage capacity above which performance degrades. Finally, we anticipate that temporal differences in excitatory and inhibitory synaptic transmission may be important to preserve NTA’s stimuli selectivity.

Our model makes several predictions. In contrast to balanced amplification, in which the network operating regime depends solely on the connectivity, an ensemble involved in NTA transitions from a non-ISN to an ISN state. This transition is consistent with noise variability observed in sensory cortices (Hennequin et al., 2018) and could be tested experimentally by probing the paradoxical effect under different stimulation conditions (Fig. 2G-H, Fig. S3). Moreover, NTA predicts that onset activity provides a better stimulus encoding and its activity is correlated with the fixed-point activity. This signature is different from purely non-normal amplification mechanisms which would involve a wave of neuronal activity across several distinct ensembles similar to a synfire chain. The difference should be clearly discernible in data. Since NTA relies on symmetric excitation between ensemble neurons, it suggests normal dynamics in which distinct ensembles first activate and then inactivate. The resulting dynamics have weak rotational components (cf. Fig. 6D) as seen in some experiments (Marshel et al., 2019). Strong non-normal amplification, on the other hand, relies on sequential activation of multiple ensembles, associated with pronounced rotational dynamics (Hennequin et al., 2014; Gillett et al., 2020), as for instance observed in motor areas (Churchland et al., 2012). Although both non-normal mechanisms and NTA are likely to co-exist in the brain, we speculate that strong NTA is best suited for, and thus most like to be found in, sensory systems.

In summary, we introduced a general theoretical framework of selective transient signal amplification in recurrent networks. Our approach derives from the minimal assumptions of a nonlinear neuronal transfer function, symmetric excitation within neuronal ensembles, and STP. Importantly, our analysis revealed the functional benefits of STP and EI co-tuning, both pervasively found in sensory circuits. Finally, our work suggests that transient onset responses rather than steady-state activity are ideally suited for coactivity-based stimulus encoding and provides several testable predictions.

## Methods

### Stability conditions for supralinear networks

The dynamics of a neuronal ensemble consisting of one excitatory and one inhibitory population with a supralinear, power law input-output function can be described as follows:

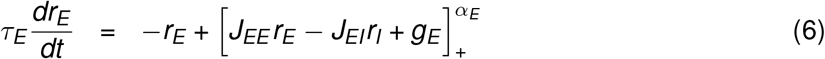

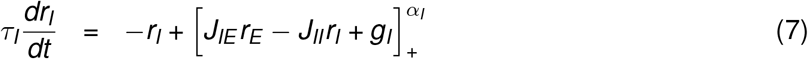

The Jacobian **M** of the system is given by

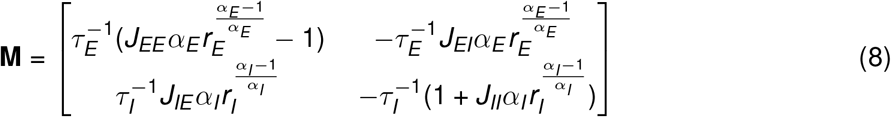

To ensure that the system is stable, the product of **M**’s eigenvalues λ_1_λ_2_, which is equivalent to the determinant of **M**, has to be positive. In addition, the sum of the two eigenvalues λ_1_ + λ_2_, which corresponds to Tr(**M**), has to be negative. We therefore obtained the following two stability conditions

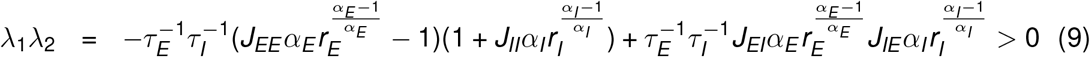

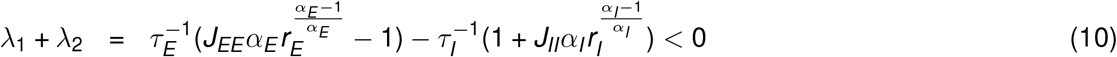

Notably, the stability conditions depend on the firing rate of the excitatory population *r_E_* and the inhibitory population *r_I_*. Since firing rates are input-dependent, the stability of supralinear networks is input-dependent. In contrast, in linear networks in which *α_E_* = *α_I_* = 1, the conditions can be simplified to

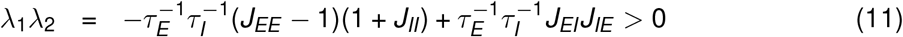

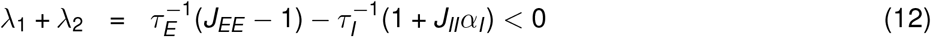

and are thus input-independent.

### ISN index for supralinear networks

If an ensemble is unstable without feedback inhibition, then the ensemble is an ISN (Tsodyks et al., 1997). To determine whether a given system is an ISN, we analyzed the stability of the E-E subnetwork, which is determined by the real part of the leading eigenvalue of the Jacobian of the E-E subnetwork. In the following we call this leading eigenvalue the “ISN index”, which is defined as follows:

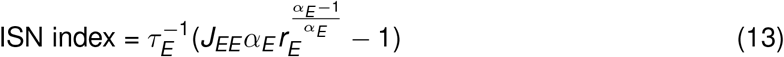

A positive ISN index indicates the system is an ISN. Otherwise, the system is non-ISN. For supralinear networks in which *α_E_* > 1, the ISN index depends on the firing rates, inputs can therefore switch the network from non-ISN to ISN. In contrast, *α_E_* = 1 for linear networks which renders the ISN index firing rate independent. And the ISN index in linear networks solely depends on the recurrent E-E connection strength *J_EE_*.

### Characteristic function

To investigate how network stability changes with input, we trace the steps of Kraynyukova and Tchumatchenko (2018) and define the characteristic function *F*(*z*) as follows:

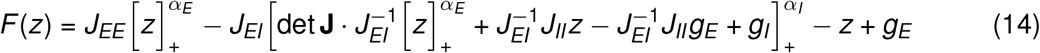

where

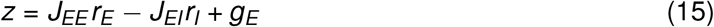

is the current into the excitatory population. The characteristic function simplifies the original two-dimensional system to a one-dimensional system, and the zero crossings of *F*(*z*) correspond to the fixed points of the original system. For *z* ≥ 0, we note:

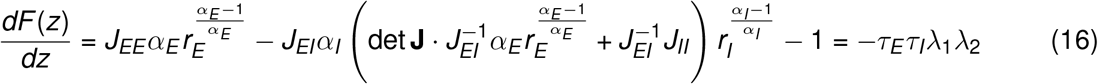

Therefore, if the derivative of *F*(*z*) evaluated at one of its roots is positive, the corresponding fixed point is a saddle point. Note that as *r_E_* and *r_I_* increase, the term in parenthesis becomes dominant. Thus, to ensure that λ_1_λ_2_ is negative also for large *r_E_* and *r_I_*, the determinant of the weight matrix det **J** has to be positive. Therefore, det **J** has a decisive impact on the curvature of *F*(*z*). In systems with negative determinant, *F*(*z*) bends upwards for large *z*. In contrast, *F*(*z*) asymptotically bends downwards in systems with positive determinant. Hence, the high-activity steady-state of systems with negative determinant is unstable. In addition, we can simplify the above condition to the determinant of the weight matrix which is a necessary condition for network stability:

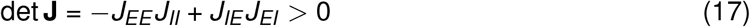

To investigate how the network stability changes with input *g_E_*, we examined how *F*(*z*) varies with changing input *g_E_* by calculating the derivative of *F*(*z*) with respect to *g_E_*,

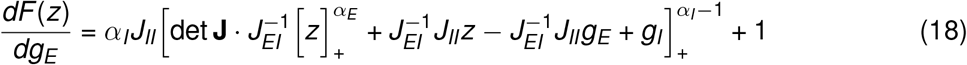

Since 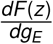 is positive, increasing *g_E_* always shifts *F*(*z*) upwards, eventually leading to the vanishing of all roots and, thus, unstable dynamics in supralinear networks with negative det **J**. In scenarios in which feedforward input to the inhibitory population also changes, we have

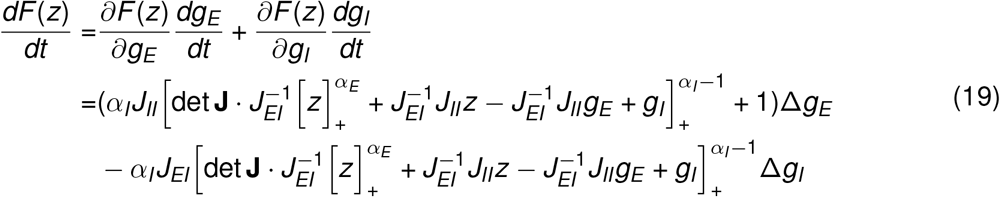

When the change in stimulation strength into the excitatory (Δ*g_E_*) and the inhibitory population (Δ*g_I_*) are the same, 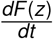 is always positive provided *J_II_* is greater than *J_EI_*. Hence, depending on the value of 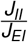, stimulation can lead to unstable network dynamics even when the input to the inhibitory population increases more than to the excitatory population.

### Spike-frequency adaptation

We modeled SFA of excitatory neurons as an activity-dependent negative feedback current (Benda and Herz, 2003; Brette and Gerstner, 2005):

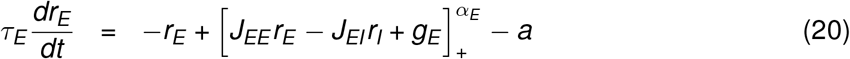

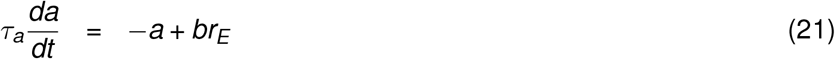

where *a* is the adaptation variable, *τ_a_* is the adaptation time constant, and *b* is the adaptation strength.

### Short-term plasticity

We modeled E-to-E STD following previous work (Tsodyks and Markram, 1997; Varela et al., 1997):

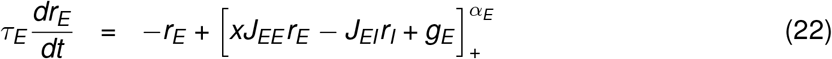

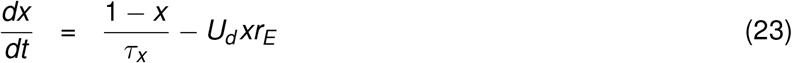

where *x* is the depression variable, which is limited to the interval (0, 1], *τ_x_* is the depression time constant, and *U_d_* is the depression rate. The steady-state solution *x*\* of STD is given by

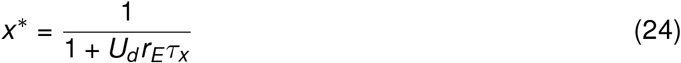

Similarly, we modeled E-to-I STF as

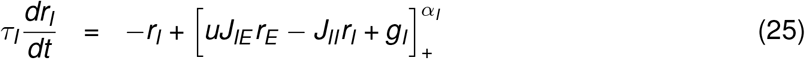

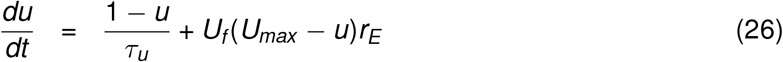

where *u* is the facilitation variable constrained to the interval [1, *U_max_*), *U_max_* is the maximal facilitation value, *τ_u_* is the time constant of STF, and *U_f_* is the facilitation rate. The steady-state solution *u*\* is given by

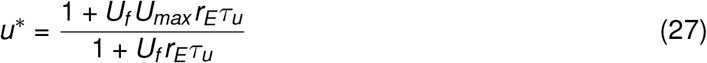

### Characteristic function with additional mechanisms

To visualize how network stability changes in the presence of SFA, we modified the characteristic function by including the adaptation strength *b*. The characteristic function with SFA is given by:

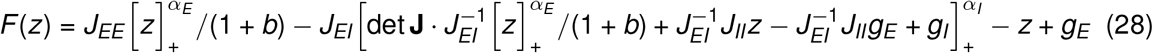

where

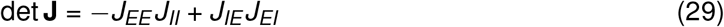

Similarly, for networks with E-to-E STD, we can modify the characteristic function by including the depression variable *x*. The characteristic function with E-to-E STD is given by:

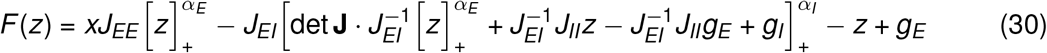

where

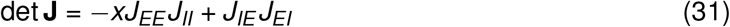

### Stability conditions for supralinear networks with additional mechanisms

To ensure network stability, Equation (10) and the simplified stability condition in Equation (17) have to be satisfied. In the presence of E-to-E STD, this results in the following conditions:

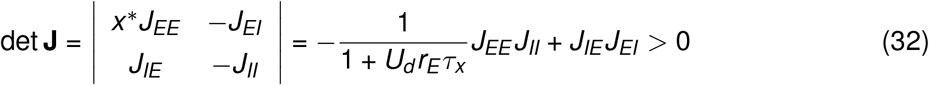

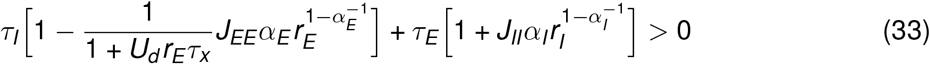

In the case of unstable dynamics, *r_E_* goes to infinity due to run-away excitation. However, the depression variable *x* approaches zero in this limit, as 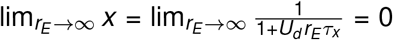. Therefore, STD terminates excitatory run-away dynamics. As a result, the ensemble transiently becomes an unstable system at stimulus onset. But assuming separation of timescales *τ_x_* ≫ *τ_E_*, STD ensures that the stability conditions are always satisfied.

In the presence of E-to-I STF, we found a set of similar conditions

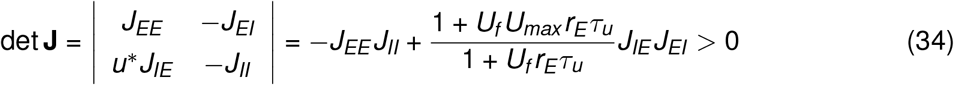

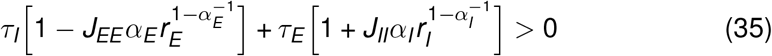

Assuming that *α_E_* = *α_I_* = *α*, the second condition becomes

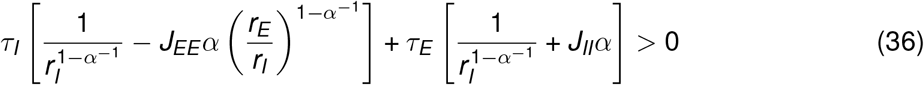

Substituting the firing rates with the current into excitatory population *z*, we then had

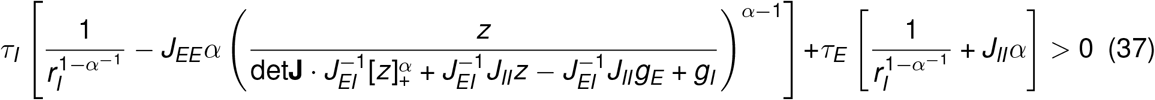

Importantly, we can guarantee that the first condition is satisfied by choosing *U_max_* sufficiently large. Since the denominator det 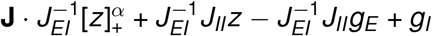 grows faster for *z* ≫ 1, the second condition is also satisfied for large *r_E_*.

In contrast, SFA does not affect the synaptic weights, the determinant of the weight matrix therefore remains negative. Although the system can have two fixed points during stimulation in the presence of weak SFA, the high-activity fixed point is always unstable. Consequently, in the presence of strong adaptation, the system exhibits oscillatory behavior (Fig. S1). To illustrate that SFA induces a limit cycle in networks that would otherwise be unstable, we considered the simplified system without the inhibitory population. The dynamics of the purely excitatory network with SFA are then given as follows:

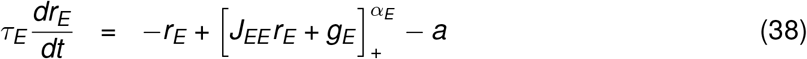

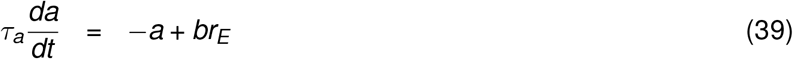

The Jacobian of the system is given as

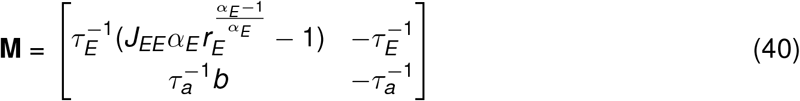

To ensure the stability of the network, the trace of the Jacobian has to be negative

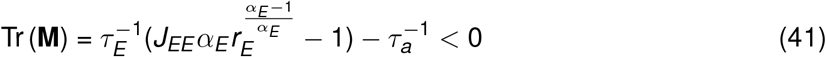

To satisfy this condition when *J_EE_* or *r_E_* are large, the time constant *τ_a_* of SFA has to be small. In addition, the determinant of the Jacobian has to be positive

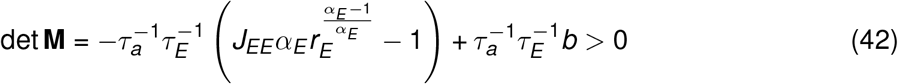

To fulfill this condition, the negative feedback controlled by *b* has to be strong. However, if both *b* and *τ_a_* are large, the determinant is positive but the trace switches from negative to positive. As a result, the system undergoes a Hopf bifurcation and exhibits oscillatory behavior (Van Vreeswijk and Hansel, 2001).

### Uni-stability conditions

The system is said to be “uni-stable”, when it has a single stable fixed point. We first identified the uni-stability condition for networks with global inhibition. To that end, we considered a general network with *N* excitatory populations and *N* inhibitory populations. To treat this problem analytically, we did not take STP into account in our analysis. The Jacobian matrix of networks with global inhibition **Q**, can be written as follows,

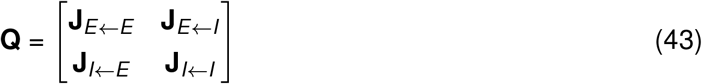

where **J**_*E←E*_, **J**_*E*←*I*_, **J**_*I*←*E*_, and **J**_*I*←*I*_ are *N* by *N* block matrices defined below.

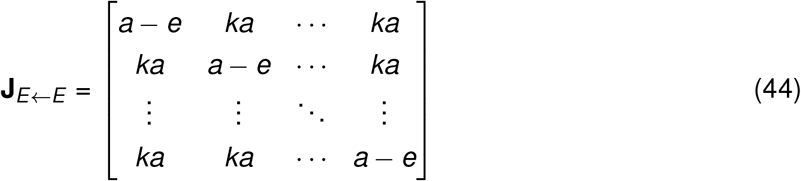

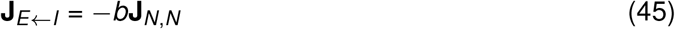

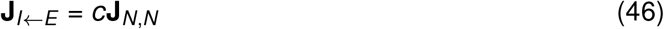

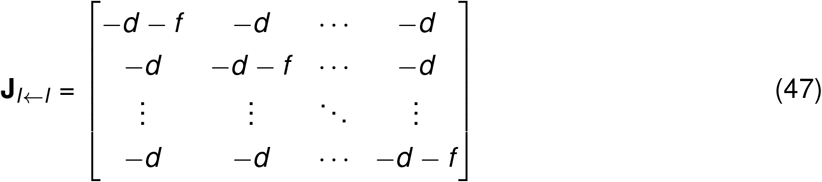

where 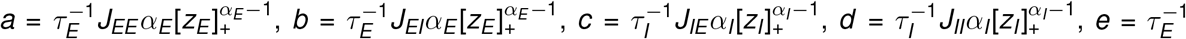, and 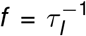. Here, *z_E_* and *z_I_* denote the total current into the excitatory and inhibitory population, respectively. Note that all these parameters are non-negative. Parameter *k* controls the excitatory connection strength across different populations. **J**_*N,N*_ is a *N* by *N* matrix of ones.

The eigenvalues of the Jacobian **Q** are roots of its characteristic polynomial,

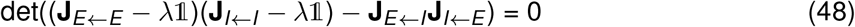

where 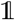 represents the identity matrix of size *N*. The characteristic polynomial can be expanded to:

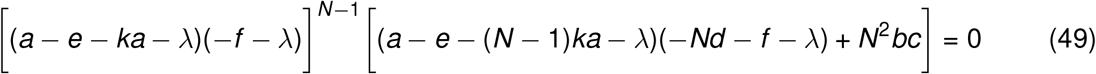

We therefore had four distinct eigenvalues:

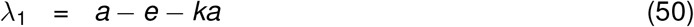

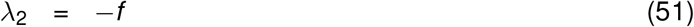

and

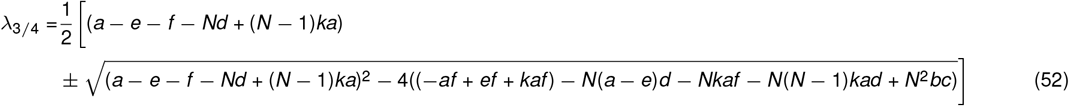

Note that the eigenvalues λ_1_ and λ_2_ have an algebraic and geometric multiplicity of (*N*-1), whereas the eigenvalues λ_3_ and λ_4_ have an algebraic and geometric multiplicity of 1.

In analogy to networks with global inhibition, the Jacobian matrix of networks with co-tuned inhibition **R**, can be written as

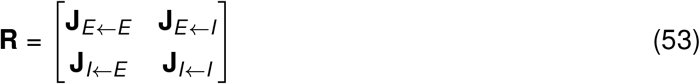

where **J**_*E*←*E*_, **J**_*E*←*I*_, **J**_*I*←*E*_, and **J**_*I*←*I*_ are *N* by *N* block matrices defined as follows:

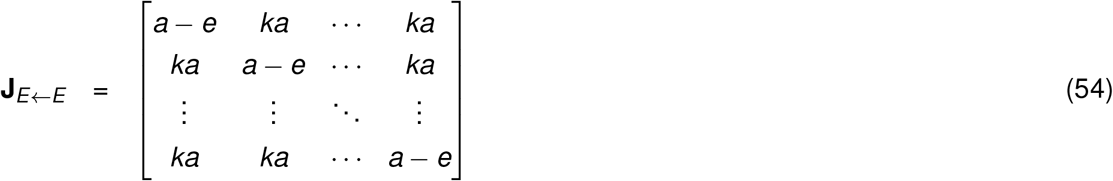

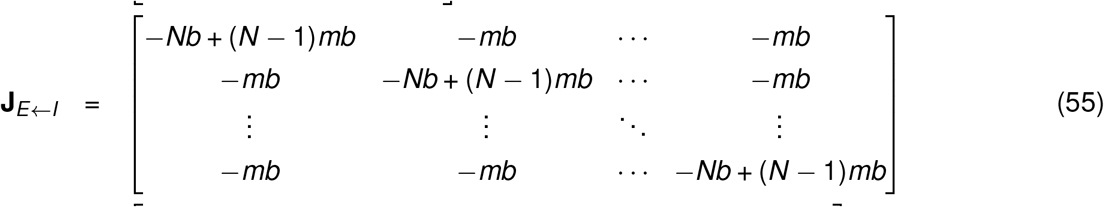

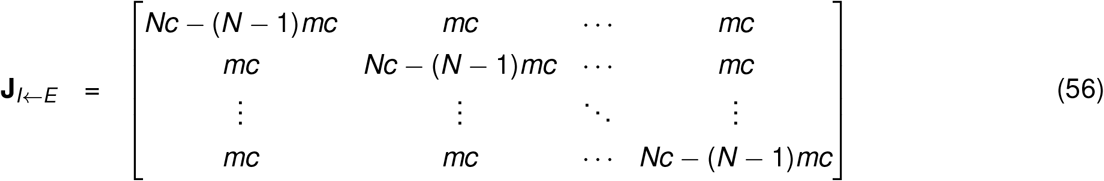

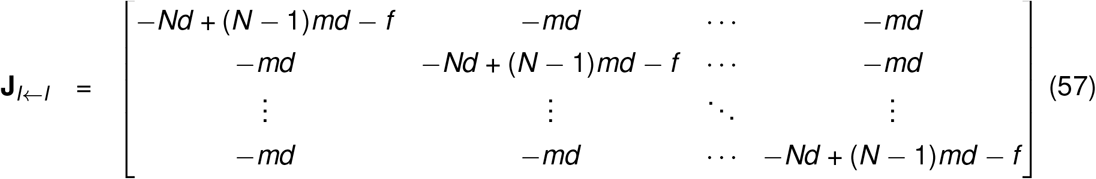

where *m* controls the degree of co-tuning in the network. If *m* = 0, the network decouples into *N* independent ensembles and inhibition is perfectly co-tuned with excitation. In the case *m* =1, inhibition is global and the block matrices become identical to the above case of global inhibition. The eigenvalues of the matrix **R** are given as the roots of the characteristic polynomial defined by:

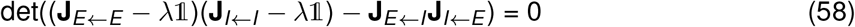

which yields the following expression:

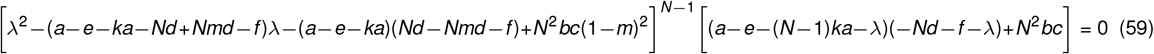

We therefore had four distinct eigenvalues:

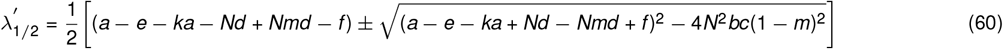

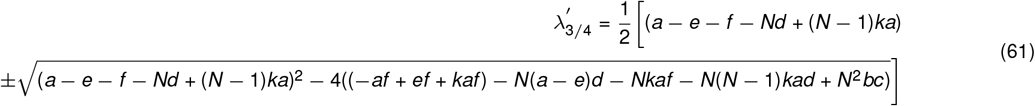

The eigenvalues 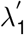 and 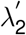 have an algebraic and geometric multiplicity of (*N*-1), whereas the eigenvalues 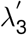 and 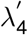 have an algebraic and geometric multiplicity of 1. We noted that 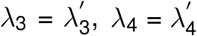.

To compare under which conditions networks with different structures are uni-stable, we examined the different eigenvalues derived above. As λ_2_ < 0, and 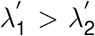, we only had to compare 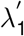 to λ_1_. For networks with co-tuned inhibition, we have *m* < 1,

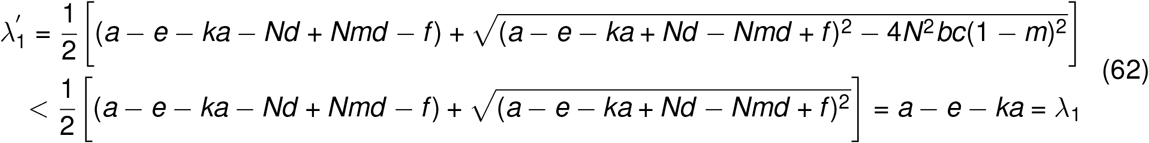

The inequality, 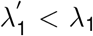, indicates that networks with co-tuned inhibition have a broad parameter regime in which they are uni-stable than networks with global inhibition. Note that in the absence of a saturating nonlinearity of the input-output function and in the absence of any additional stabilization mechanisms, systems with positive eigenvalues of the Jacobian are unstable. In this case, networks with co-tuned inhibition have a broad parameter regime of being stable than networks with global inhibition.

To visualize the conditions in a two-dimensional plane, we reduced the conditions into a function of *a* and *d*. For Fig. 3C, *k* = 0.1, *m* = 0.5 and *bc* = 0.9*ad*.

### Distance to the decision boundary

To calculate the distance to the decision boundary in Fig. 4, Fig. 5, Fig. S6 and Fig. S7, we first projected the excitatory activity in Phase 2 onto a two-dimensional Cartesian coordinate system in which the *x* axis is the activity of the first excitatory ensemble *r*_*E*1_ and the *y* axis is the activity of the second excitatory ensemble *r*_*E*2_. We then computed the distance between the projected data and the decision boundary which corresponds to the diagonal line in the coordinate system.

### Inhibitory feedback pathways for suppressing unwanted neural activation

To identify the important neural pathways for the suppression of unwanted neural activation, we analyzed how the activity of the second excitatory ensemble *r*_*E*2_ changes with the input to the first excitatory ensemble *g*_*E*1_. To that end, we considered a general weight matrix for networks with two interacting ensembles

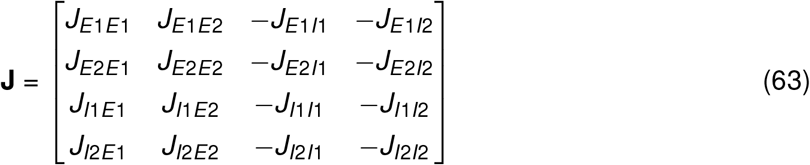

We can write the change in firing rate of the excitatory population in the second ensemble *δr*_*E*2_ as a function of the change in the input to the other *δg*_*E*1_:

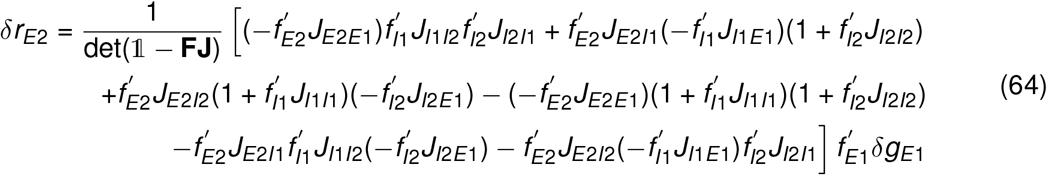

where 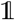 is the identity matrix. And **F** is given by

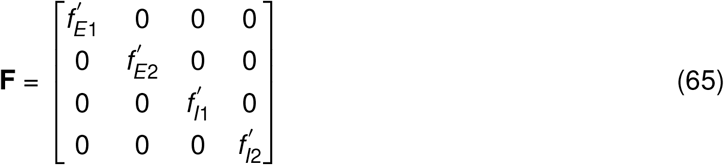

where 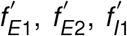 and 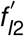 are the derivatives of the input-output functions evaluated at the fixed point.

Assuming that 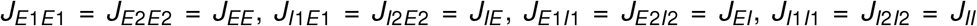, 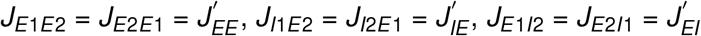 and 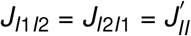, we find

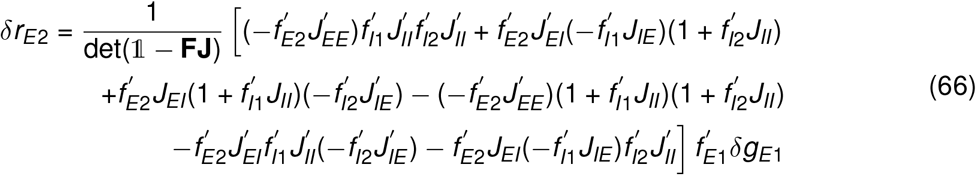

By further assuming that the weight strengths across ensembles are weak and ignoring the corresponding higher-order terms, we get

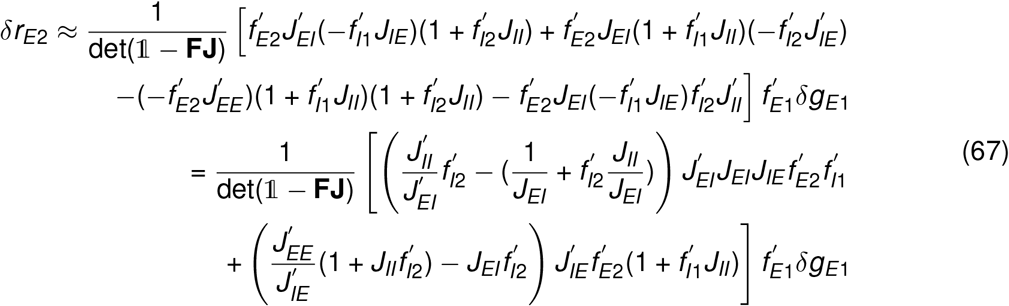

Note that 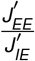 and 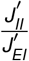 are terms regulating the respective excitatory and inhibitory input from one ensemble to the excitatory and inhibitory population in another ensemble. The term 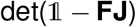 is positive to ensure the stability of the system.

To suppress the activity of the excitatory population in the second ensemble *r*_*E*2_, in other words, to ensure that *δr*_*E*2_ < 0, 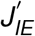 or/and 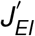 have to be large. Therefore, we identified 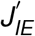 and 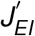 as important synaptic connections which lead to suppression of the unwanted neural activation, suggesting that inhibition can be provided via 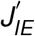 through the *E*1-*I*2-*E*2 pathway or via 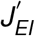 through the *E*1-*I*1-*E*2 pathway.

For Fig. 4–5, the rate-based model consists of two ensembles, each of which is composed of 100 excitatory and 25 inhibitory neurons with all-to-all connectivity.

### Spiking neural network model

The spiking neural network model was composed of *N_E_* excitatory and *N_I_* inhibitory leaky integrate-and-fire neurons. Neurons were randomly connected with probability of 20%. The dynamics of membrane potential of neuron *i*, *U_i_*, are given by (Zenke et al., 2015):

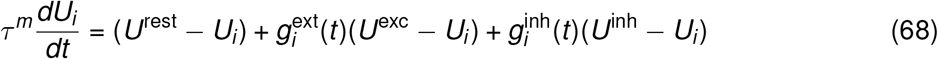

Here, *τ^m^* is the membrane time constant and *U*^rest^ is the resting potential. Spikes are triggered when the membrane potential reaches the spiking threshold *U*^thr^. After a spike is emitted, the membrane potential is reset to *U*^rest^ and the neuron enters a refractory period of *τ*^ref^. Inhibitory neurons obeyed the same integrate-and-fire formalism but with a shorter membrane time constant.

Excitatory synapses contain a fast AMPA component and a slow NMDA component. The dynamics of the excitatory conductance are described by:

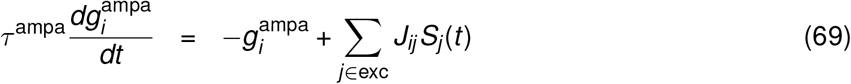

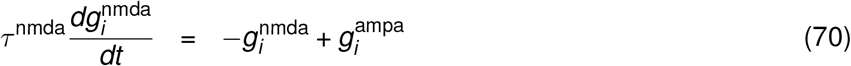

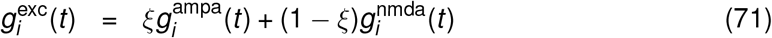

Here, *J_ij_* denotes the synaptic strength from neuron *j* to neuron *i*. If the connection does not exist, *J_ij_* was set to 0. *S_j_*(*t*) is the spike train of neuron *j*, which is defined as 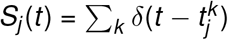, where *δ* is the Dirac delta function and 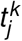 the spikes times *k* of neuron *j*. *ξ* is a weighting parameter. The dynamics of inhibitory conductances are governed by:

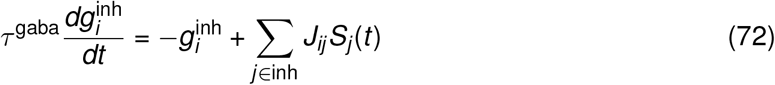

In the spiking neural network models, SFA of excitatory neurons is modeled as follows,

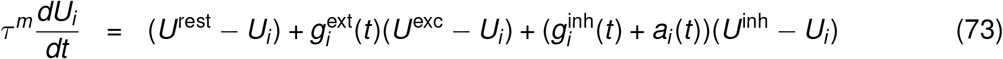

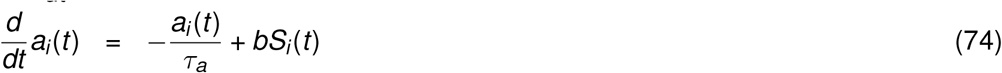

where *i* is the index of excitatory neurons.

The dynamics of E-to-E STD are given by

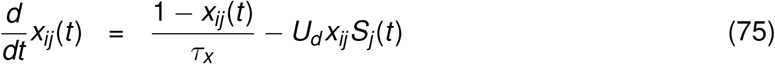

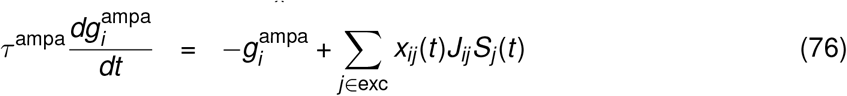

where *i* represents the index of excitatory neurons.

The dynamics of E-to-I STF are governed by

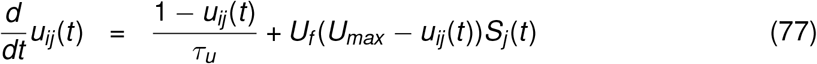

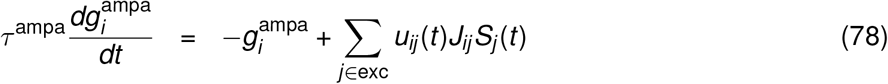

where *i* denotes the index of inhibitory neurons.

For Fig. 6, each excitatory and inhibitory neuron received external excitatory input from 300 neurons firing with Poisson statistics at an average firing rate of 0.1 Hz at baseline. During stimulation, the excitatory neurons corresponding to the activated pattern received external excitatory input from 300 neurons firing with Poisson statistics at an average firing rate of 0.5 Hz. The pattern activity with each stored pattern is quantified by the dot product of the neural activity with the stored pattern. And neural activity is computed by the instantaneous firing rates with 10 ms bin size. The difference in pattern activity for the peak amplitude is calculated by subtracting the average maximal pattern activity of the unstimulated patterns from the maximal pattern activity of the activated pattern. Similarly, the difference in pattern activity for the fixed point is calculated by subtracting the average pattern activity of the unstimulated patterns at the fixed point from the pattern activity of the activated pattern at the fixed point. Fixed point activity is computed by averaging the activity of the middle 1 second within the 2-second stimulation period.

For Fig. S5, each excitatory and inhibitory neuron received external excitatory input from 300 neurons firing with Poisson statistics at an average firing rate of 0.1 Hz at the baseline. During stimulation, each excitatory neuron received external excitatory input from 300 neurons firing with Poisson statistics at an average firing rate of 0.3 Hz.

### Simulations

Simulations were performed in Python and Mathematica. All differential equations were implemented by Euler integration with a time step of 0.1 ms. All simulation parameters are listed in Tables 1–7. The simulation source code to reproduce the figures is publicly available at https://github.com/fmi-basel/gzenke-nonlinear-transient-amplification.

**Table 1:** Parameters for Figure 1C-E.

| Table 1: Parameters for Figure 1C-E |  |  |  |
| --- | --- | --- | --- |
| Symbol | Value | Unit | Description |
| $J_{EE}$ | 1.8 | - | E-to-E connection strength |
| $J_{IE}$ | 1.0 | - | E-to-I connection strength |
| $J_{EI}$ | 1.0 | - | I-to-E connection strength |
| $J_{II}$ | 0.6 | - | I-to-I connection strength |
| $\alpha_E$ | 2 | - | power of excitatory input-output function |
| $\alpha_I$ | 2 | - | power of inhibitory input-output function |
| $\tau_E$ | 20 | ms | time constant of excitatory firing dynamics |
| $\tau_I$ | 10 | ms | time constant of inhibitory firing dynamics |
| $g_E^{bs}$ | 1.55 | - | input to the $E$ population at baseline |
| $g_E^{stim}$ | 3.0 | - | input to the $E$ population during stimulation |
| $g_I$ | 2.0 | - | input to the $I$ population |
| Parameters for Figure 1F |  |  |  |
| $J_{IE}$ | 0.45 | - | E-to-I connection strength |
| $J_{EI}$ | 1.0 | - | I-to-E connection strength |
| $J_{II}$ | 1.5 | - | I-to-I connection strength |

**Table 2:** Parameters for Figure 2.

| Table 2: Parameters for Figure 2 |  |  |  |
| --- | --- | --- | --- |
| Symbol | Value | Unit | Description |
| $\tau_a$ | 200 | ms | time constant of SFA |
| $b$ | 1.0 | - | strength of SFA |
| $\tau_x$ | 200 | ms | time constant of STD |
| $U_d$ | 1.0 | - | depression rate |
| $\tau_u$ | 200 | ms | time constant of STF |
| $U_f$ | 1.0 | - | facilitation rate |
| $U_{max}$ | 6.0 | - | maximal facilitation value |

Note that these values are also applied elsewhere unless mentioned otherwise.

**Table 3:** Parameters for Figure 3 bi/multi-stable example.

| Table 3: Parameters for Figure 3 bi/multi-stable example |  |  |  |
| --- | --- | --- | --- |
| Symbol | Value | Unit | Description |
| $J_{EE}$ | 1.4 | - | within-ensemble E-to-E connection strength |
| $J_{IE}$ | 0.6 | - | within-ensemble E-to-I connection strength |
| $J_{EI}$ | 1.0 | - | within-ensemble I-to-E connection strength |
| $J_{II}$ | 0.6 | - | within-ensemble I-to-I connection strength |
| $J'_{EE}$ | 0.14 | - | inter-ensemble E-to-E connection strength |
| $J'_{IE}$ | 0.6 | - | inter-ensemble E-to-I connection strength |
| $J'_{EI}$ | 1.0 | - | inter-ensemble I-to-E connection strength |
| $J'_{II}$ | 0.6 | - | inter-ensemble I-to-I connection strength |
| $g_{E1}^{bs}$ | 2.2 | - | input to the $E1$ population at baseline |
| $g_{E1}^{stim}$ | 3.0 | - | input to the $E1$ population during stimulation |
| $g_{E2}$ | 2.2 | - | input to the $E2$ population |
| $g_I$ | 2.0 | - | input to the $I$ population |

| Parameters for Figure 3 uni-stable example |  |  |  |
| --- | --- | --- | --- |
| $J_{EE}$ | 1.3 | - | within-ensemble E-to-E connection strength |
| $J'_{EE}$ | 0.13 | - | inter-ensemble E-to-E connection strength |

**Table 4:** Parameters for Figure 4–5.

| Symbol | Value | Unit | Description |
| --- | --- | --- | --- |
| $N_E$ | 200 | - | number of excitatory neurons |
| $N_I$ | 50 | - | number of inhibitory neurons |
| $N$ | 2 | - | number of ensembles |
| $J_{EE}$ | $1.2/(N_E/2 - 1)$ | - | within-ensemble E-to-E connection strength |
| $J_{IE}$ | $1.0/(N_E/2)$ | - | within-ensemble E-to-I connection strength |
| $J_{EI}$ | $1.0/(N_I/2)$ | - | within-ensemble I-to-E connection strength |
| $J_{II}$ | $1.0/(N_I/2 - 1)$ | - | within-ensemble I-to-I connection strength |
| $J'_{EE}$ | $0.36/(N_E/2 - 1)$ | - | inter-ensemble E-to-E connection strength |
| $J'_{IE}$ | $0.4/(N_E/2)$ | - | inter-ensemble E-to-I connection strength |
| $J'_{EI}$ | $0.1/(N_I/2)$ | - | inter-ensemble I-to-E connection strength |
| $J'_{II}$ | $0.1/(N_I/2)$ | - | inter-ensemble I-to-I connection strength |
| $g_I$ | 2.0 | - | input to the $I$ population |
| **Parameters for Figure 4** |  |  |  |
| $g_{E1}^{bs}$ | 1.35 | - | input to the $E1$ population at baseline |
| $g_{E1}^{stim}$ | 4.0 | - | input to the $E1$ population during stimulation |
| $g_{E2}$ | 1.35 | - | input to the $E2$ population |
| **Parameters for Figure 5** |  |  |  |
| $g_{E1}^{bs}$ | 1.35 | - | input to the $E1$ population at baseline |
| $g_{E1}^{stim}$ | $1.35 + (4.0 - 1.35)(1-p)$ | - | input to the $E1$ population during stimulation |
| $g_{E2}^{bs}$ | 1.35 | - | input to the $E2$ population at baseline |
| $g_{E2}^{stim}$ | $1.35 + (4.0 - 1.35)p$ | - | input to the $E2$ population during stimulation |

Here, $p$ is a parameter between 0 and 1 controlling the additional inputs to $E1$ and $E2$ .

**Table 5:** Parameters for Figure 6.

| Symbol | Value | Unit | Description |
| --- | --- | --- | --- |
| $N_E$ | 400 | - | number of excitatory neurons |
| $N_I$ | 100 | - | number of inhibitory neurons |
| $U^{\text{rest}}$ | -70 | mV | resting membrane potential |
| $U^{\text{exc}}$ | 0 | mV | excitatory reversal potential |
| $U^{\text{inh}}$ | -80 | mV | inhibitory reversal potential |
| $\tau^{\text{ref}}$ | 3 | ms | duration of refractory period |
| $\tau_{\text{exc}}^m$ | 20 | ms | membrane time constant of excitatory neurons |
| $\tau_{\text{inh}}^m$ | 10 | ms | membrane time constant of inhibitory neurons |
| $\tau_{\text{ampa}}$ | 5 | ms | time constant of AMPA receptor |
| $\tau_{\text{gaba}}$ | 10 | ms | time constant of GABA receptor |
| $\tau_{\text{nmda}}$ | 100 | ms | time constant of NMDA receptor |
| $\xi$ | 0.5 | - | receptor weighting factor |
| $J_{EE}$ | 0.19 | - | within-ensemble E-to-E connection strength |
| $J_{IE}$ | 0.10 | - | within-ensemble E-to-I connection strength |
| $J_{EI}$ | 0.10 | - | within-ensemble I-to-E connection strength |
| $J_{II}$ | 0.06 | - | within-ensemble I-to-I connection strength |
| $J'_{EE}$ | 0.019 | - | inter-ensemble E-to-E connection strength |
| $J'_{IE}$ | 0.05 | - | inter-ensemble E-to-I connection strength |
| $J'_{EI}$ | 0.04 | - | inter-ensemble I-to-E connection strength |
| $J'_{II}$ | 0.006 | - | inter-ensemble I-to-I connection strength |

**Table 6:** Parameters for Figure S5.

| Symbol | Value | Unit | Description |
| --- | --- | --- | --- |
| $N_E$ | 400 | - | number of excitatory neurons |
| $N_I$ | 100 | - | number of inhibitory neurons |
| $J_{EE}$ | 0.05 | - | E-to-E connection strength |
| $J_{IE}$ | 0.02 | - | E-to-I connection strength |
| $J_{EI}$ | 0.05 | - | I-to-E connection strength |
| $J_{II}$ | 0.03 | - | I-to-I connection strength |

**Table 7:** Parameters for Figure S9.

| Symbol | Value | Unit | Description |
| --- | --- | --- | --- |
| $J_{EE}$ | 0.5 | - | E-to-E connection strength |
| $J_{IE}$ | 0.45 | - | E-to-I connection strength |
| $J_{EI}$ | 1.0 | - | I-to-E connection strength |
| $J_{II}$ | 1.5 | - | I-to-I connection strength |
| $g_E^{bs}$ | 0.5 | - | input to the $E$ population at baseline |
| $g_I^{bs}$ | 1.5 | - | input to the $I$ population at baseline |

## Acknowledgments

We thank Rainer W. Friedrich, Claire Meissner-Bernard, William F. Podlaski, and members of the Zenke Group for comments and discussions. This work was supported by the Novartis Research Foundation.

## Author contributions

Y.K.W. and F.Z. conceived the study. Y.K.W. performed model analyses and simulations. Y.K.W. and F.Z. wrote the manuscript.

## Competing interests

The authors declare no competing interests.

## Supplementary Figures

**Fig. S1.**
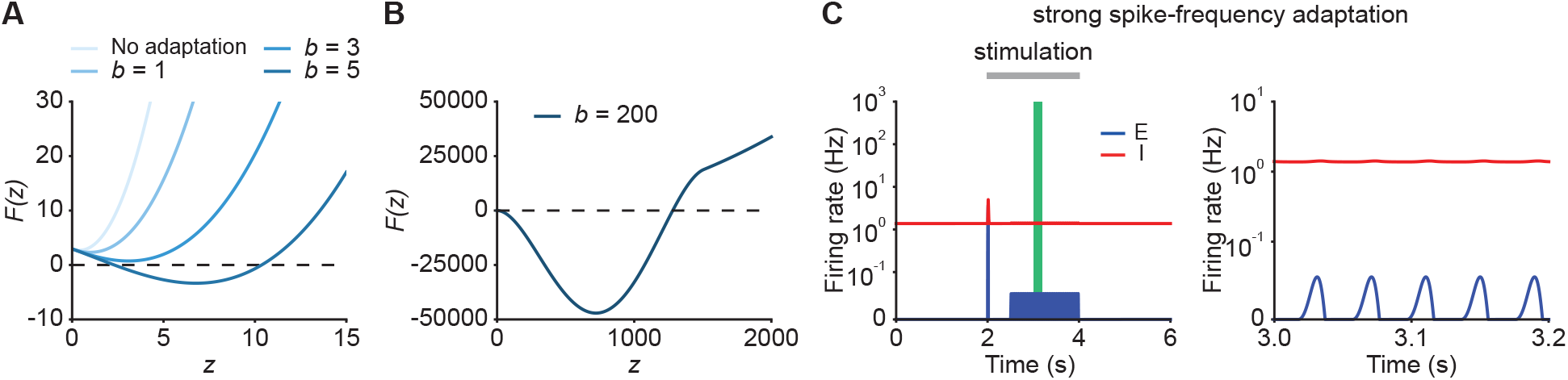
**(A)** Characteristic function F(z) in networks with weak SFA which cannot stabilize run-away activity. Different colors represent different adaptation strengths *b*. **(B)** Characteristic function F(z) in networks with strong SFA capable of generating a limit cycle. **(C)** Firing rates of the excitatory (blue) and inhibitory population (red) in the presence of strong SFA (left). The zoomed-in activity from 3.0 s to 3.2 s (right) corresponding to the green period (left) indicates oscillatory behavior in networks with strong SFA.

**Fig. S2.**
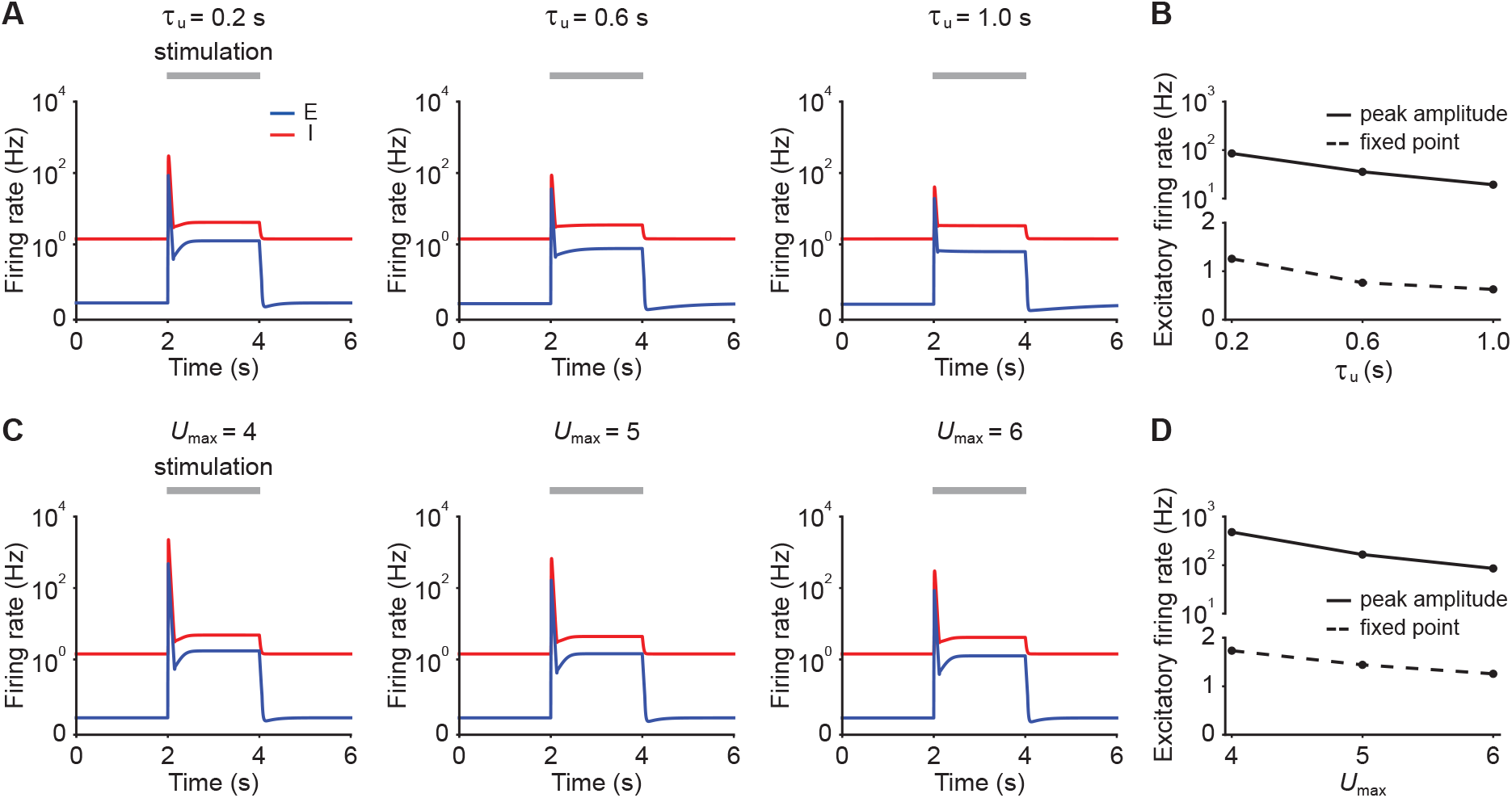
**(A)** Firing rates of the excitatory (blue) and inhibitory population (red) in response to stimulation in the presence of E-to-I STF with different time constants *τ_u_*. The stimulation period from 2s to 4s is marked with the gray bar. The stimulation is implemented by changing input *g_E_*. **(B)** Peak amplitude (solid line) and fixed-point activity (dashed line) of the excitatory population during stimulation with different STF time constants. **(C and D)** Same as A and B, but with different maximum allowed facilitation levels *U_max_*.

**Fig. S3.**
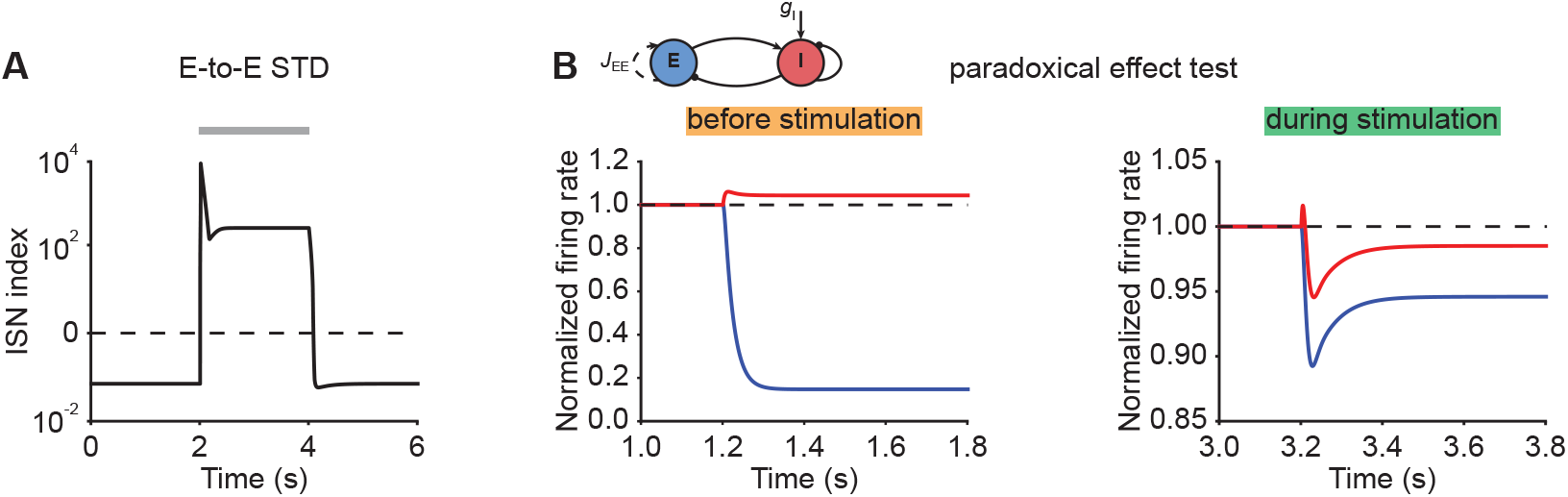
**(A)** ISN index of the network with E-to-E STD in Fig. 2B (left). The horizontal dashed line indicates an ISN index of 0. **(B)** The normalized firing rates of the excitatory (blue) and inhibitory population (red) when injecting excitatory input into the inhibitory population of an active ensemble (cf. Fig. 2B) starting at 1.2 s while the ensemble receives external stimulation (left), and at 3.2 s (right). The horizontal dashed lines indicate a normalized firing rate of 1.0.

**Fig. S4.**
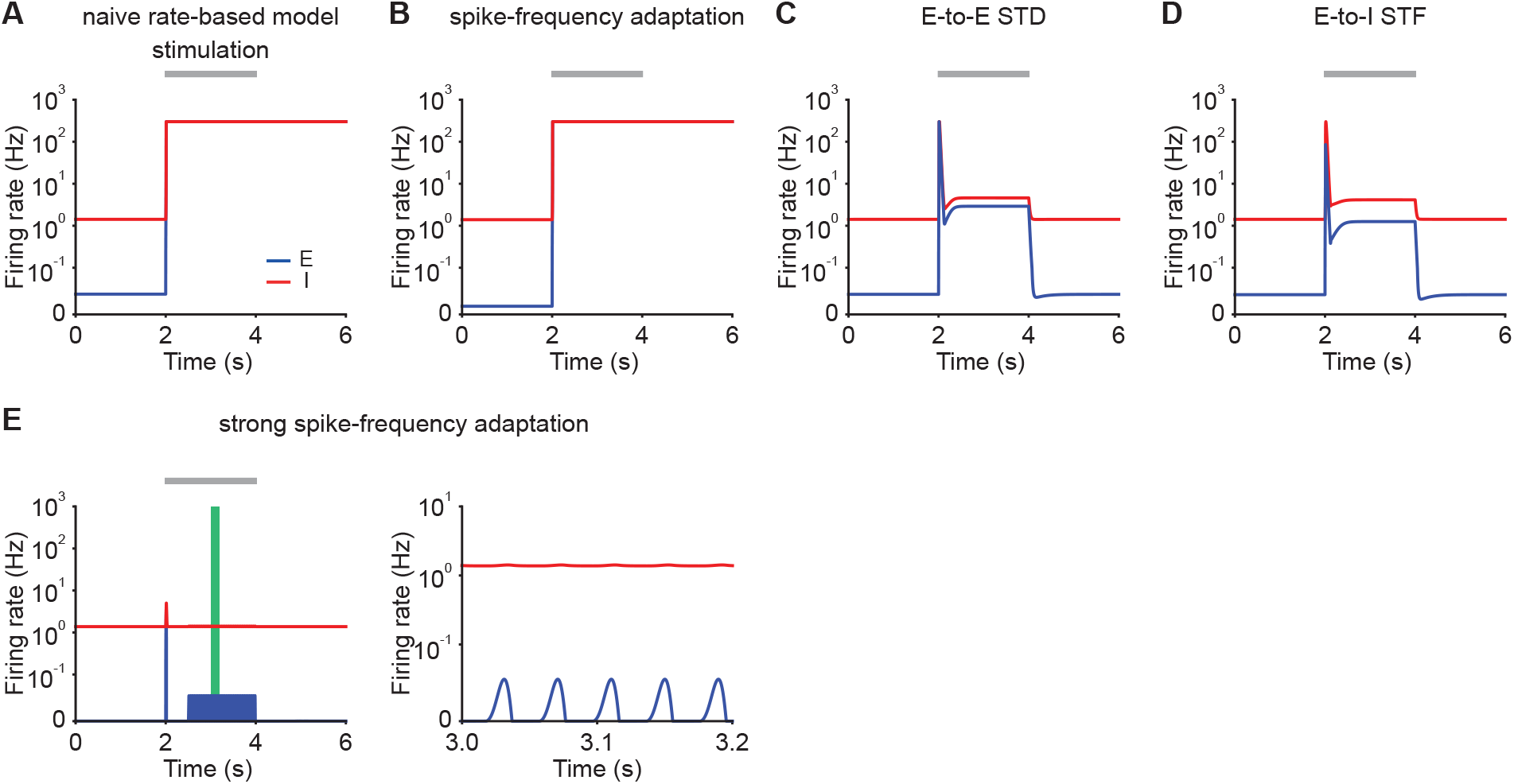
**(A)** Firing rates of the excitatory (blue) and inhibitory population (red) in response to stimulation in a rate-based model. Additional excitatory inputs are injected into excitatory neurons in the period marked in gray. Firing rates are capped at 300 Hz. **(B)** Same as A but incorporating SFA. **(C)** Same as A but incorporating E-to-E STD. **(D)** Same as A but incorporating E-to-I STF. **(E)** Same as B but incorporating stronger SFA (left). The zoomed-in activity from 3.0 s to 3.2 s (right) corresponding to the green period (left) indicates oscillatory behavior in networks with strong SFA.

**Fig. S5.**
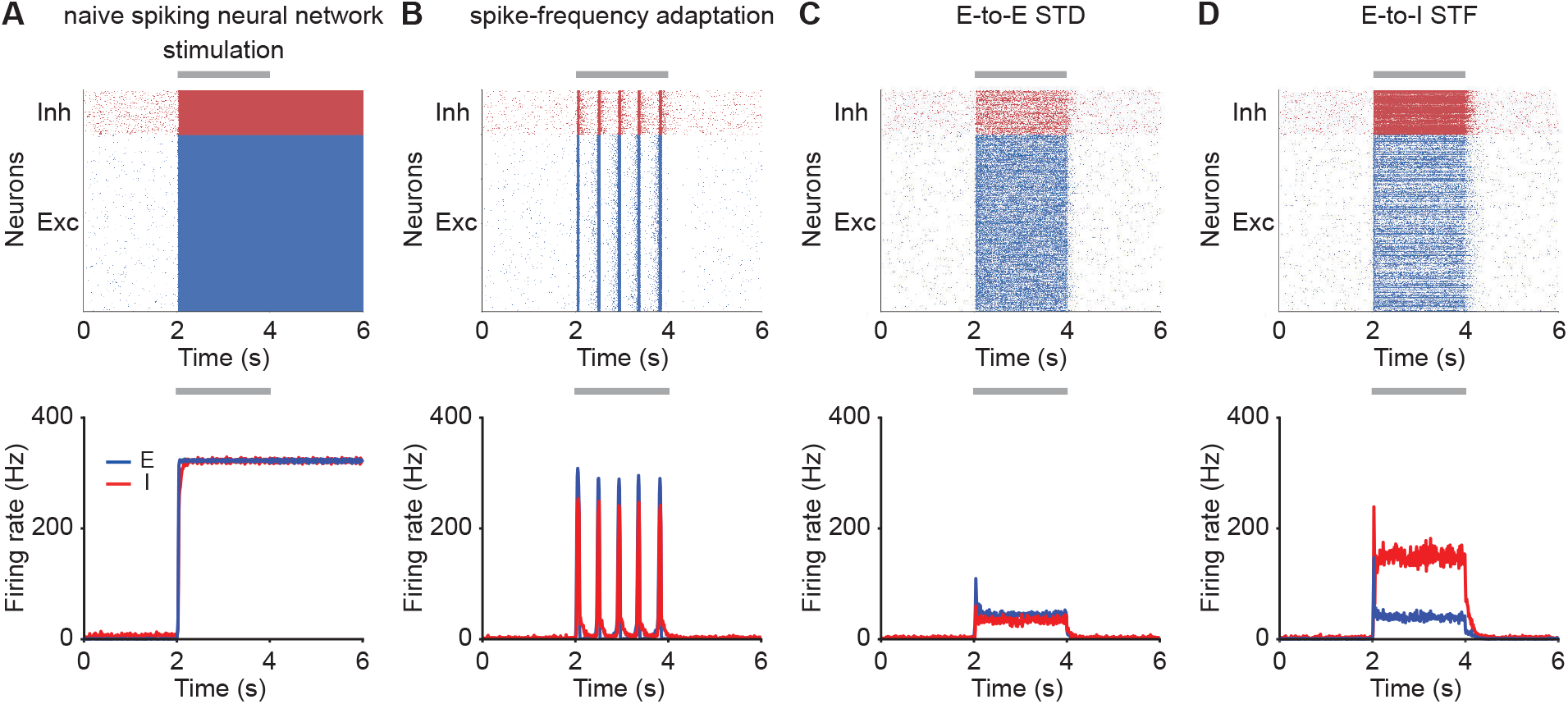
**(A)** Spike raster plot of the excitatory (top, blue) and inhibitory population (top, red) in response to stimulation in a spiking neural network model, and firing rates calculated with 10 ms time bins (bottom). Additional excitatory inputs are injected into excitatory neurons in the period marked in gray. **(B)** Same as A but incorporating SFA. **(C)** Same as A but incorporating E-to-E STD. **(D)** Same as A but incorporating E-to-I STF.

**Fig. S6.**
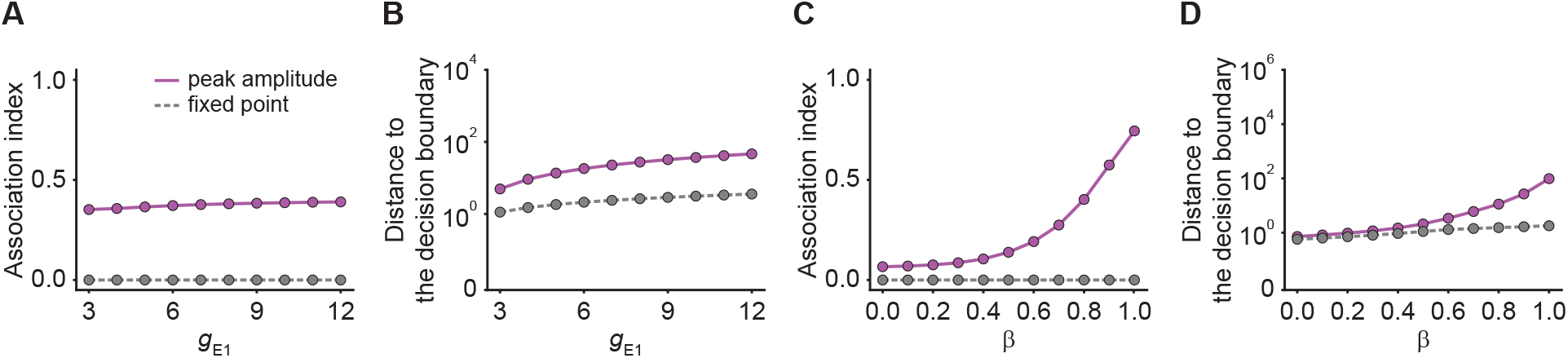
**(A)** Association index as a function of input *g*_*E*1_ for the onset peak amplitude (magenta solid line) and fixed-point activity (gray dashed line) for E-to-I STF. **(B)** Distance to the decision boundary as a function of input *g*_*E*1_ for the onset peak amplitude (magenta solid line) and fixed-point activity (gray dashed line) for E-to-I STF. **(C and D)** Same as A and B but as a function of *β*, which controls the inner- and inter-ensemble connection strength.

**Fig. S7.**
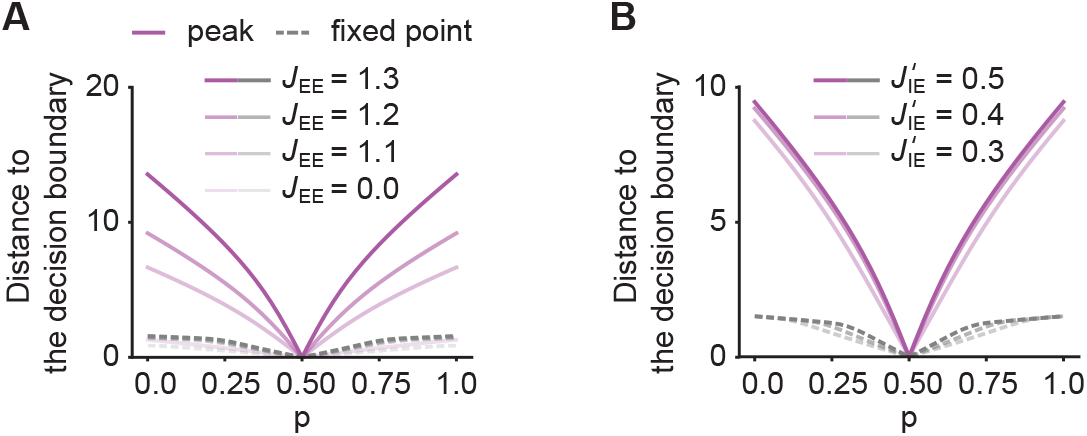
**(A)** Distance to the decision boundary as a function of *p* for the onset peak amplitude (magenta solid lines) and fixed-point activity (gray dashed lines) for E-to-I STF. Different levels of brightness represent different recurrent E-to-E connection strengths *J_EE_*. **(B)** Same as A but with different E-to-I connection across ensembles strengths 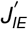.

**Fig. S8.**
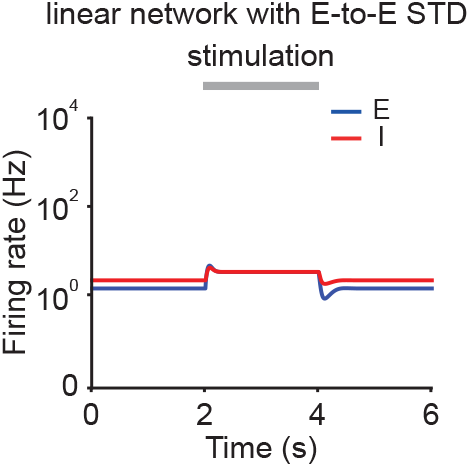
Firing rates of the excitatory (blue) and inhibitory population (red) in linear networks with E-to-E STD. During stimulation (gray bar) additional input is injected into the excitatory population. Same as Fig. 2B (left) but with *α_E_* = *α_I_* = 1.

**Fig. S9.**
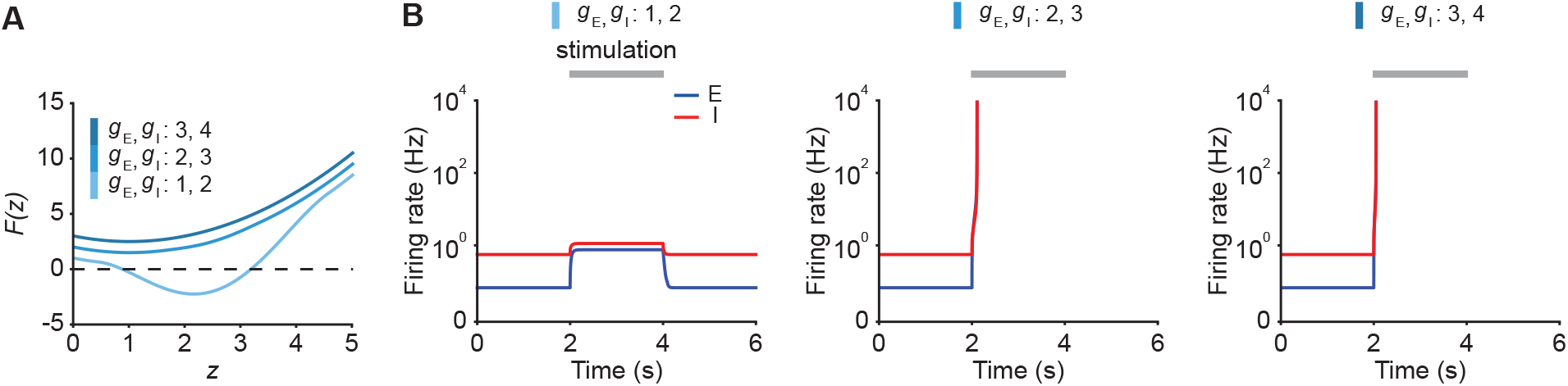
**(A)** Characteristic function *F*(*z*) for different inputs *g_E_* and *g_I_*. **(B)** Firing rates of the excitatory (blue) and inhibitory population (red) in response to stimulation from 2–4s (gray bar). During stimulation *g_E_* and *g_I_* are simultaneously changed to the stated values.

## Notes

### Competing Interest Statement

The authors have declared no competing interest.

